# Interpreting tree ensemble machine learning models with endoR

**DOI:** 10.1101/2022.01.03.474763

**Authors:** Albane Ruaud, Niklas Pfister, Ruth E Ley, Nicholas D Youngblut

## Abstract

Tree ensemble machine learning models are increasingly used in microbiome science as they are compatible with the compositional, high-dimensional, and sparse structure of sequence-based microbiome data. While such models are often good at predicting phenotypes based on microbiome data, they only yield limited insights into how microbial taxa or genomic content may be associated. Results: We developed endoR, a method to interpret a fitted tree ensemble model. First, endoR simplifies the fitted model into a decision ensemble from which it then extracts information on the importance of individual features and their pairwise interactions and also visualizes these data as an interpretable network. Both the network and importance scores derived from endoR provide insights into how features, and interactions between them, contribute to the predictive performance of the fitted model. Adjustable regularization and bootstrapping help reduce the complexity and ensure that only essential parts of the model are retained. We assessed the performance of endoR on both simulated and real metagenomic data. We found endoR to infer true associations with more or comparable accuracy than other commonly used approaches while easing and enhancing model interpretation. Using endoR, we also confirmed published results on gut microbiome differences between cirrhotic and healthy individuals. Finally, we utilized endoR to gain insights into components of the microbiome that predict the presence of human gut methanogens, as these hydrogen-consumers are expected to interact with fermenting bacteria in a complex syntrophic network. Specifically, we analyzed a global metagenome dataset of 2203 individuals and confirmed the previously reported association between *Methanobacteriaceae* and *Christensenellales*. Additionally, we observed that *Methanobacteriaceae* are associated with a network of hydrogen-producing bacteria. Conclusion: Our method accurately captures how tree ensembles use features and interactions between them to predict a response. As demonstrated by our applications, the resultant visualizations and summary outputs facilitate model interpretation and enable the generation of novel hypotheses about complex systems. An implementation of endoR is available as an open-source R-package on GitHub (https://github.com/leylabmpi/endoR).

---

The gut microbiome plays critical roles in many aspects of human physiology, such as digestion, immunity, and development (1–3), and has been implicated in a number of diseases (4) such as inflammatory bowel disease (IBD) (5, 6), obesity (7, 8), diabetes (9), and cancer (10, 11). The low cost of fecal microbiome sequencing allows researchers and clinicians to relate disease states to microbiome data and also to investigate possible microbial involvement in disease (12–14).

Machine Learning (ML) models have been shown to accurately predict human host phenotypes from gut microbiome taxonomic and genomic data (15–18). While the complexity of these models can capture interactions between variables in such data, it also complicates their interpretation. This consequently limits insights into relationships between the microbiome and human characteristics. Random forest (RF) models (19), a type of tree ensemble model, often achieve the best accuracies for predictions made with microbiome data (15– 18). A RF consists of a combination of decision trees. Each partitions all observations into subsamples with similar response values, based on a set of features. For example, a decision tree may show that diseased individuals generally have high abundances of microbes A and B, but low abundances of microbe C. Hundreds of decision trees are built from random subsamples of features and observations and aggregated to make predictions. This procedure is called bootstrap aggregation or bagging (19); it generally leads to high accuracies with less overfitting but increases model complexity (20).

The model complexity can be mitigated by reducing the number of features via feature selection (FS): the pre-selection of relevant features to include in the final model (21–24). These pre-selection approaches are often based on different measures of how important a specific feature is for the prediction. Such measures are also the state-of-the-art for model interpretation. The most common feature importances are the Gini and permutation importances (19, 25), though many others exist (26–30). Recently developed FS and feature importance methods are gaining popularity within microbiome science. For instance, Ai et al. (26) utilized FS by mutual information to identify specific microbes predictive of colorectal cancers. Alternatively, Gou et al. (31) used SHapley Additive exPlanations (SHAP, Lundberg and Lee (28)) to select microbiome features associated with type 2 diabetes that they then correlated to host genetics and risk factors using generalized linear models.

Shapley values measure the contribution of variables to the prediction of each observation (32) and can be estimated through various methods (33). For instance, the SHAP method additively decomposes predictions into separate parts corresponding to each variable (28). As Shapley values generate local, per-observation interpretations, they generally do not address the question of the global associations of features with the response (33). Furthermore, SHAP makes the assumption that variables have additive effects, although tree ensembles are not additive models, therefore resulting in potentially biased estimates of feature interactions (34). Finally, SHAP interactions are calculated only for pairs of variables, rendering their interpretation challenging for high-dimensional data sets (35).

Individual decision trees can inform on variable interactions associated with predictions. Variables belonging to a same tree branch are used in concert to make predictions; they are thus more likely to be jointly associated with the response compared to variables never appearing in the same branch (36). However, as tree ensembles are generated with a greedy procedure, especially RFs, unimportant variables may occur along decision paths. To remove noise and facilitate the interpretation of tree ensembles, Friedman and Popescu (37) propose to remove unimportant variables from decision paths via lasso regression and thus create surrogate models to tree ensembles. The inTrees R-package (38) and random intersection tree algorithm (27) implement similar ideas of simplifying tree ensembles to obtain a reduced set of decisions from a forest. However, they lack the tools to interpret the simplified decisions further. Conversely, the randomForest-Explainer R-package (39) measures variable interactions by counting the number of co-occurrences of features in decision trees. However, noise is not removed from tree ensembles before measuring variable co-occurrences. The package also does not generate easy to interpret results for models fitted on high-dimensional data.

To better interpret fitted tree ensemble models, we developed endoR, a framework for interpreting tree ensemble models. endoR utilizes decisions extracted from fitted models to infer associations between features and measures their contribution to the decision ensemble (Figure 1). The endoR workflow consists of extracting all decisions from a tree ensemble model, simplifying them, and then calculating the importance and influence of variables and interactions among pairs of variables. More specifically, the importance measures the gain in predictive accuracy attributed to a variable (or an interaction among a pair of variables), while the influence measures how the inclusion of a variable (or an interaction among a pair of variables) changes model predictions. Results are displayed as multiple intelligible plots to enhance the readability of feature and interaction importances and influences.

**Figure 1.**
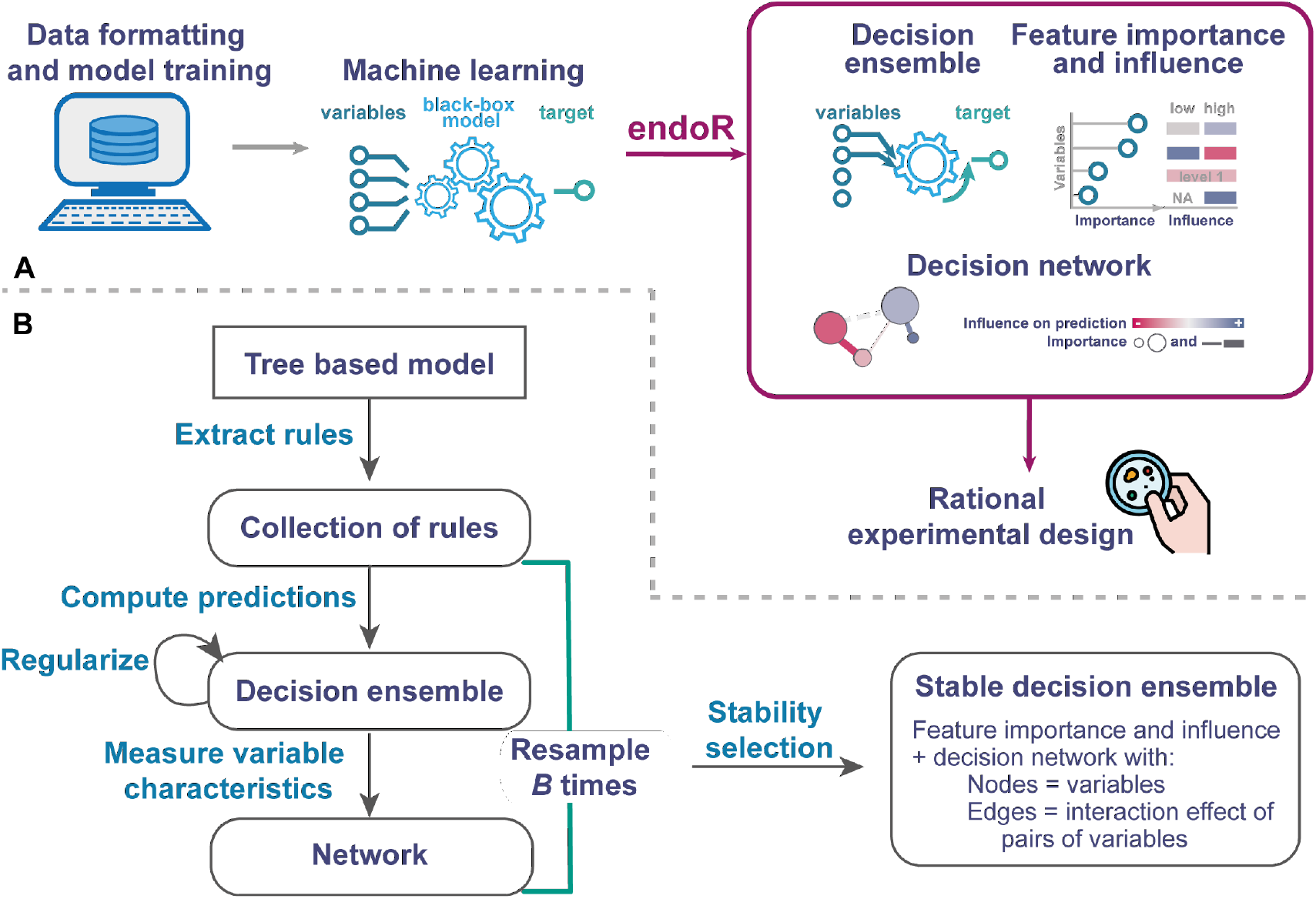
Description of the endoR method workflow. A/ General overview of the workflow from data acquisition to the visualization of a network. endoR is applied to a trained classification or regression tree ensemble model. The model is first simplified into a decision ensemble, which is used to calculate the feature importance and influence on predictions. The resultant metrics are displayed in a summary plot listing the feature importance and influence, and as a decision network. The decision network illustrates the association between the response and single or pairs of variables, in regards to feature importance and influence. If the influence of a variable depends on other variables, it will be visible in the network via edges between these nodes. B/ Steps taken by endoR to generate a stable network. endoR accepts tree ensemble models that were made with the XGBoost, gbm, randomForest or ranger R-packages (43–46). Regularization is optional and consists of simplifying decisions and the decision ensemble to reduce noise. The procedure can be repeated on *B* bootstraps to select stable decisions prior to constructing the final network.

Notably, endoR generates a decision network which visualizes the fitted model as follows: (i) nodes represent the features used in the model, with their size and color encoding the feature importance and influence (i.e., the strength and direction of association with the response, respectively); (ii) edges represent interaction effects on the response between two features. Similarly, the width and color encodes the interaction importance and influence, respectively.

We benchmarked endoR on both a fully simulated data set and a real metagenome data set (40), both with an artificially generated phenotype as response. In particular, we compared endoR with state-of-the-art procedures commonly used for analyzing microbiome data. Altogether, our results showed that endoR successfully extracts complex interactions from tree ensemble models and performs better or comparable to existing methods. We then employed endoR on a metagenome dataset published by Qin et al. (41), in which the original study identified certain gut microbiome features to be associated with cirrhosis. From a single application of endoR, we were able to recover all major results of the original study and expand upon them by identifying additional oral bacteria colonizing the gut of patients with cirrhosis and the depletion of bacteria associated with healthy microbiome (42). Finally, we used endoR to explore patterns of gut microbial relative abundances predictive of the presence of *Methanobacteriaceae* in human guts. The presence of these methanogens was strongly associated with the presence of members of the *CAG-138* family (order *Christensenellales*), specifically with the *Phil-1* genus, as well as with members of the *Oscillospirales* order. Moreover, host traits such as the body mass index (BMI), were not predictive of the presence of *Methanobacteriaceae*, suggesting that the microbiome composition primarily determines *Methanobacteriaceae* prevalence across human populations. Taken together, the application of endoR provides new perspectives on the prevalence of *Methanobacteriaceae* in the human gut and their plausible interactions with members of the gut microbiome.

## Results

### Interpreting tree ensemble models with endoR

Tree ensemble models are often used in microbiome science, although their interpretation is limited by their complexity. endoR overcomes this issue by taking a fitted model as input and visualizing the most relevant parts of the model in a feature importance and influence plot and a decision network. It is implemented as an R-package and accepts fitted RF and gradient boosted tree models (both for regression and classification tasks) generated using the XGBoost, gbm, randomForest, or ranger R-packages (43–46). We illustrate the use of endoR on a data set consisting of 2147 human gut metagenomes, with relative abundances of 520 taxa (including species, genus, and family taxonomic rank), and an artificial phenotype. The phenotype was a binary response variable taking the values ‘-1’ or ‘1’, it was simulated using 9 randomly selected taxa and a randomly generated categorical variable separating samples in 4 groups (labelled a, b, c, and d). Selected taxa could be from the species, genus, or family taxonomic levels to mirror the range of interactions that can occur in reality between between microbial clades of varying taxonomic resolution. The mechanism generating the artificial phenotype, which we use as response variable in the classification below, is detailed in Methods and Table 2; the taxa associated with the artificial phenotype within each group are visualized in Figure 2 A-F. Our goal is to recover as much information about the mechanism generating the artificial phenotype as possible. For example, is it possible to determine that in Group a, high relative abundances of *Alistipes A*, and high relative abundances of *Marvinbryantia sp900066075* lead to a positive value of the artificial phenotype? Even though Figure 2 A-F may suggest that this is an easy classification task, it is in fact highly non-trivial; the simulation involves high order interactions (up to order 4) in a high-dimensional setting (521 variables) with strong dependencies between the features from real metagenomes. Tree ensemble models such as RFs excel in these settings, but they do not provide methods for extracting complex information about the model. This is the gap that endoR aims to fill.

**Figure 2.**
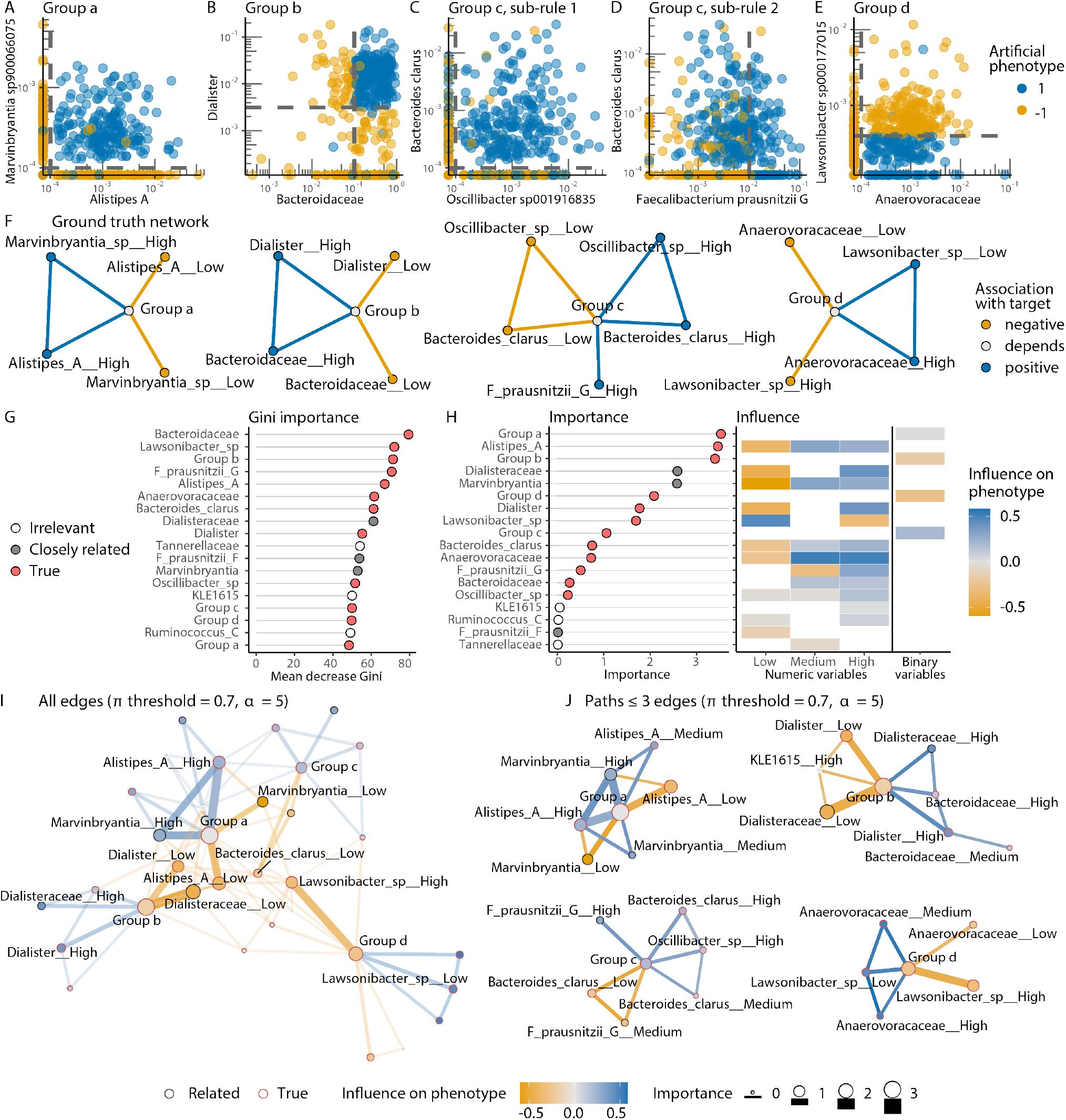
endoR captures interactions predictive of an artificial phenotype from a random forest fitted on real metagenomes. A-E/ Real metagenomes with an artificial phenotype (AP): samples were separated into 4 groups (labelled a-d), a binary response variable (‘1’ = blue, ‘-1’ = yellow) was simulated so that it could be predicted from a set of decisions based on the ‘group’ categorical feature and specific, randomly chosen microbial abundance features (e.g., ‘Alistipes A’). Dashed grey lines denote thresholds in the predetermined decisions used to make the response variable and are described in Table 2 (e.g., the response variable is ‘1’ if samples belong to Group a and have non-null relative abundances of both *Alistipes A* and *Marvinbryantia sp900066075*). For samples in Group c, the response variable was built with an ‘OR’ rule (i.e., ‘Group = c & ((B. clarus *>* 0 & Oscillibacter sp001916835 *>* 0) | F. prausnitzii G *>* 10^*−*2^)’), so each of the two sub-rules are shown in C/ and D/. F/ Ground truth network of features derived from the response variable generation procedure described in A/. Pairs of variables predicting ‘1’ are linked by a blue edge (‘positive’) and those predicting ‘-1’ by a yellow edge (‘negative’). Variables for which high values are predictive of ‘1’ have a blue node color (‘positive’) and a yellow node color if high values are predictive of ‘-1’ (‘negative’). If high values are predictive of ‘1’ or ‘-1’ depending of other variable values (e.g., Group b predicts ‘1’ if V3 takes high values, but ‘-1’ if V3 has low values), the color is grey (‘depends’). G-H/ Variable importances from the RF model as measured by the mean decrease in Gini impurity and endoR. The point color indicates whether the features were used to construct the response (‘True’) and those taxonomically related to them (‘closely related’), with ‘closely related’ defined as the immediate parent or child taxonomic classification in the taxonomy hierarchy (e.g., the *Bacteroides* genus is the child of the *Bacteroidaceae* family, while *Bacteroidaceae* is the parent of *Bacteroides*). I/ Full decision network extracted by endoR from a RF model trained on the dataset described in A/. Only the 20 features with the highest feature importance are labelled. The edge transparency is inversely proportional to the importance for I/ only. J/ Same network as shown in I/, but edges with lowest interaction importance were removed to obtain paths between nodes of length ≤ 3. All features are labelled.

First, we fit a model that predicts the artificial phenotype from the taxa (i.e., the relative abundances of species, genera, and families) and the ‘group’ variable. Given the high number of features, FS was used before fitting a RF (see Methods). The fitted model in this case has a cross-validation (CV) generalization error of 85.19±2.36 for the accuracy and 0.70±0.05 for Cohen’s *κ*. Next, we apply endoR, which outputs two plots: an importance and influence plot (Figure 2 H) and a decision network (Figure 2 I-J).

The importance and influence plot shows the feature importance and influence for a single variable (Figure 2 H). The importance measures how much a single variable improves the overall prediction of the model; it is similar to other well-established importance measures such as the Gini importance (given in Figure 2 G) but is more accurate in its ranking of variables, such that irrelevant taxa were given the lowest feature importance by endoR but not by the Gini importance (Figure 2 G and H). As shown in our simulations below, the endoR feature importance improves on standard importance measures for tree ensemble models. As a complement, the influence measures the change in predicted value due to the variable. For binary features, it indicates whether samples falling in that category take on average higher or lower response variable values. For instance, Figure 2 H shows that samples from Group d are more likely to have a ‘-1’ artificial phenotype (orange) while there is no clear association for samples in Group a (grey). Hence, Group a is important for predictions, but its association with the artificial phenotype may mostly depend on other features. For numeric features (taxa), the influence similarly shows how the variable is associated with the response in the final decision ensemble. To help readability, numeric variables are split into levels defined by value ranges. The number of levels is pre-specified by the user; here, ‘low’, ‘medium’, and ‘high’ values of each variable are assessed. If a level does not appear in the decision ensemble, its influence cannot be calculated, and so it is left blank in the plot. Figure 2 H shows that ‘low’ relative abundances of *Alistipes A* are associated with the ‘-1’ phenotype while ‘medium’ and ‘high’ relative abundances of this taxon are associated with the ‘1’ phenotype. The importance and influence plot thus provides an overview of important features and how they affect the response on average.

The decision network allows for a more detailed analysis. The nodes in the network correspond to each possible value of categorical features (e.g., ‘Group a’) and levels of numerical features (the level is indicated by ‘ Level’, e.g., ‘Marvinbryantia High’). The size of the node corresponds to the importance, while the color encodes the influence. Edges correspond to interaction effects, while size and color indicate importance and influence of the interaction, respectively. Either the full network can be displayed (Figure 2 I) or only the most important paths composed by less than three edges (Figure 2 J). For example, in Figure 2 J, we can see that the network indeed separates the 4 groups into separate components and also captures a pattern specific to Group a: high relative abundances of both *Alistipes A* and *Marvinbryantia* are associated with a positive phenotype for samples in Group a. Although species *Marvinbryantia sp900066075* was the true predictor in our simulation, the genus *Marvinbryantia* was instead selected by the predictive model – likely due to the high redundancy among these closely related features. This example illustrates how endoR can depend on the fitted model. If endoR is fitted on a decision ensemble directly obtained from the true mechanism generating, rather than by fitting a predictive model, it indeed recovers the ground truth one (Supplementary Figure 1 A-C).

### Evaluation of endoR on simulated data

In this section, we summarize our findings from evaluating endoR on multiple simulated datasets; further details and additional evaluations can be found in the Supplementary Results. The evaluation is based on two simulation configurations. The first configuration, referred to as *fully simulated data* (FSD). Unlike our previous simulation used for demonstrating endoR, the FSD were constructed by simulating both the features and response variable. Features are independent from each other, normally distributed, and all predictive associations are known (illustrated in Supplementary Figure 2). The second configuration, referred to as *artificial phenotypes* (AP), is similar to the simulation used to demonstrate endoR, in that AP simulations also comprise features from published human gut metagenomes comprising 2147 samples (40) and a response variable constructed from combinations of relative abundances of randomly chosen taxa (Figure 2). Hence, predictive variables are dependent and not all predictive associations are known. A more detailed description of the data is given in the Methods section.

#### endoR is robust to changes in hyperparameters

We generated 100 FSD and 50 AP datasets, processed them with varying endoR hyperparameters, and evaluated how these changes affected the ability of endoR to recover correct edges in the decision networks (Figure 3 A and D and Supplementary Figure 5 C, F, and G-J).

**Figure 3.**
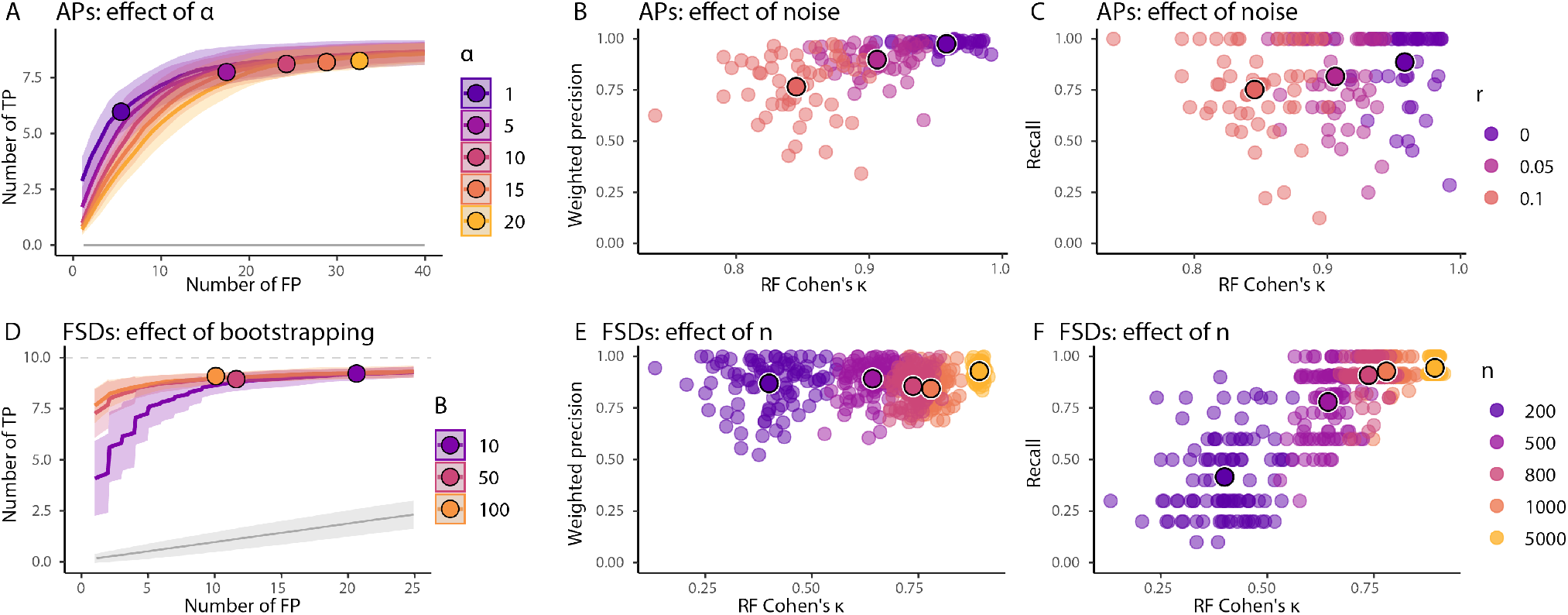
endoR’s performance is robust to hyperparameters and depends on input model. Simulation results based on 100 FSDs with *n* = 1000 observations (except when varied in E-F) and 50 APs using all observations (A-C). In all experiments, the noise was *r* = 0.05 (except when varied in B-C) and endoR was applied to fitted RFs with *α* = 5 (except when varied in A) and *B* = 10 (except when varied in D). For each dataset and parameter setting we fitted a RF and applied endoR. Then, we computed the following three metrics: Cohen’s *κ* of the RF, weighted precision and recall values of the selected edges in the stable decision ensemble, and TP/FP-curves based on the probabilities of being selected in the stable decision ensemble (see Methods). A and D/ TP/FP-curves are averaged across all datasets for a fixed parameter setting (line) and standard deviation (shaded area) are displayed. The average number of TPs and FPs expected for a randomization null model and standard deviations, are shown in grey. Large points indicate the average number of TPs and FPs in the stable ensembles generated by endoR. B-C and E-F/ Each point corresponds to the precision/recall of endoR applied to a single dataset and parameter setting. The larger traced points are the averages across all datasets for a fixed parameter setting. A/ Increasing *α* increases both the TPs and FPs. Small values of *α* effectively control the FPs without strongly impacting the recovered TPs. D/ Larger values of *B* are slightly better but endoR performs well even for small values of *B*. B-C and E-F/ As expected decreasing the noise or increasing the number of observations improves the performance of endoR both in terms of precision and recall. Importantly, there is a strong dependence of endoR performance on the performance of the fitted RF and endoR. Moreover, endoR has a good precision even for small sample sizes.

First, we explored the effect of *α*, which by construction is supposed to control the expected number of wrong decisions selected by endoR after bootstrapping. Accordingly, the number of TP and FP edges identified by endoR also increased with increasing *α* (Figure 3 A-B). Even for small values of *α*, endoR recovered many TPs while controlling for low numbers of FPs. Furthermore, regardless of *α*, TP edges were attributed the highest importances, hence resulting in high weighted precision of final decision ensembles (Supplementary Table 1).

Second, we varied the number of bootstraps on which our stability selection procedure is applied. Varying the number of bootstrap resamples between 10, 50, and 100 for FSDs, and 10 and 90 for APs, slightly increased the precision and sensitivity of endoR (Figure 3 D and Supplementary Figure 5 F), and higher bootstrap numbers decreased the overfitting of results (Supplementary Figure 3). Generally a higher value for *B* is preferable but the size is limited by computing resources (see Supplementary Figure 8 E and F for an evaluation of computing resources required). Our empirical results suggest that a value between 10 and 100 is often sufficient.

Lastly, we assessed whether endoR’s performance was affected by the method used for discretizing numeric value (e.g., grouping into ‘Low’ and ‘High’ numeric values; Supplementary Figure 4). endoR is indeed robust to the discretization procedure (Supplementary Results and Supplementary Figure 5 G-J).

#### endoR improves with the accuracy of fitted models

Since endoR interprets tree ensemble models, we evaluated the influence of the accuracy of RF models on the accuracy of endoR. We fitted RFs to 100 FSD and 50 AP datasets and applied endoR to the models. Model accuracy was altered by varying (i) noise levels via the *r* parameter (Figure 3 B-C and Supplementary Figure 5 A-B), (ii) the number of samples used to fit models (Figure 3 E-F),and (iii) the model complexity through the number of trees in the forest (Supplementary Figure 5 D-E). Model accuracy increased with higher numbers of trees, lower noise, or higher number of samples (Figure 3 B-C and E-F and Supplementary Figure 5 A-B and D-E).

On average, the weighted precision of endoR was high, even for low predictive model performances (i.e., small Cohen’s *κ*; Figure 3 E), and it increased with RF model performance as noise in data declined (Figure 3 B). Importantly, even for small sample size (e.g., *n* = 200) endoR had high weighted precision values (Figure 3 E). We attribute this to the regularization and resampling steps used in endoR, that effectively reduce the risk of overfitting. The endoR recall always increased with higher RF predictive performance (Figure 3 C and F). Furthermore, the variance of the recall across datasets decreased with increased predictive performances, meaning that although endoR produces precise networks, the probability of recovering as many true interactions as possible increases with the predictive model accuracy. Taken together, the results consistently showed that the performance of endoR depends on the quality of the input model (Figure 3 B and E).

#### endoR outperforms state-of-the-art methods for metagenome data analysis

We utilized 50 AP datasets to evaluate the performance of endoR relative to the state-of-the-art (Figure 4 and Supplementary Methods). Our evaluations included non-parametric statistical Wilcoxon rank-sum and *χ*^2^ tests, sparse covariance matrices computed with the sparCC (47) and graphical lasso (gLASSO, Friedman et al. (48)) methods, the Gini importance (19, 25), and SHAP values (28). In particular, we used RFs to extract Gini importances, SHAP values of single variables, and endoR feature and interaction importances. Given that SHAP interaction values are not readily available for RF models in R, we fitted gradient boosted models using the xgboost R-package (46) to extract SHAP values and interaction values, Gini importances, and endoR feature and interaction importances. Wilcoxon rank-sum and *χ*^2^ tests identified single variables significantly associated with the artificial phenotypes, while sparse covariance matrices discriminated pairs of variables significantly correlated in one but not the other phenotypic group.

**Figure 4.**
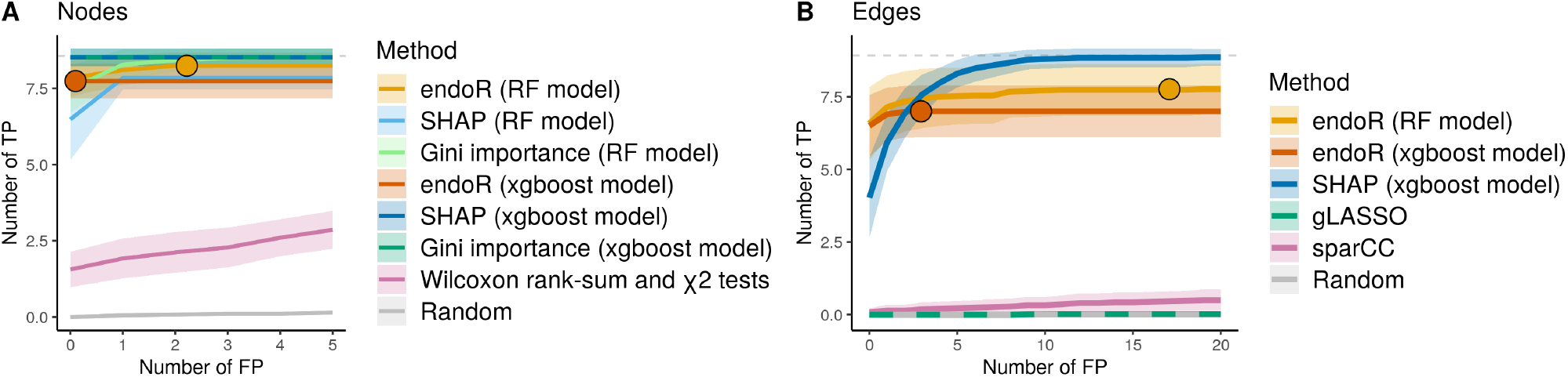
endoR is better or as good as state-of-the-art methods at identifying true variables and pairs of variables predictive of artificial phenotypes. Average (line) and standard deviation (area) of the number of identified true positive (TP) for a given number of false positive (FP). The average numbers of TP and FP in the endoR final decision ensemble are indicated with points. A/ corresponds to single variables and B/ to pairs of variables across 50 replicates of artificial phenotypes. Dashed grey lines denote the ground truth number of TP. ‘Random’ signifies results expected with a randomization null model. A/ ‘SHAP (xgboost model)’ and ‘Gini (xgboost model)’ lines are dashed due to their overlap. B/ ‘gLASSO’ and ‘Random’ lines are dashed due to their overlap. A-B/ All methods based on fitted predictive models almost perfectly ranked TP because of the FS step in model fitting. B/ endoR better discriminated TP from FP edges than SHAP. Only endoR does not return all features and interactions, hence limiting the number of FPs in the final decision ensembles, although resulting in lower recall too.

Each of the 50 APs simulated from real metagenomes was processed with all methods. Single variables and pairs of variables were ranked by the output parameters of each method and compared with the ground truth network to build TP/FP curves (Supplementary Methods). Each curve displays the number of TP variables, or interaction effects between pairs of variables, found by each method for a given number of FP on average across the 50 APs (Figure 4).

All methods that did not use a predictive model (i.e., non-parametric statistical tests, sparCC, and gLASSO), performed poorly, with accuracies nearly equivalent to random guessing (Figure 4). The generally good performance of methods based on classifiers was due to the FS step performed with gRRF (21) during model training (Figure 4). Overall, single variables were very well ranked by endoR, SHAP, and Gini importances, with close to all TP attributable to the highest importances before any FP (Figure 4 A). SHAP and Gini had a better recall, but endoR was the only method to return a subset of variables, hence limiting the number of FP.

Interactions were identified with a higher recall by endoR from RF than XGBoost models in these simulations (Figure 4 B), even though the average Cohen’s *κ* of XGBoost models was higher compared to RFs (on average across the 50 repetitions, from the mean across 10 CV sets for each replicate, Cohen’s *κ* = 0.97±0.00 and 0.91±0.03 for XG-Boost and RF models, respectively). endoR was more accurate than SHAP at ranking of interactions. Again, SHAP recall was higher but the number of FP was limited in the endoR decision ensemble due to the selection of variables via regularization. Furthermore, endoR could extract interaction importances from RF models, while SHAP is not available in R for this purpose (Figure 4 B). Hence, endoR outperformed other methods in terms of accuracy of results.

We note that the summary plots generated by endoR, specifically the feature importance and influence plot and the decision network, enable rapid assessment of important variables and the direction of their association with the response variable, as well as interactions between variables. On the contrary, SHAP values are designed to inform at the per-observation level and are not suited to provide general overview of results, especially as *p* increases (Supplementary Figures 6 and 7). Therefore, in terms of data interpretability, endoR better suits analyses of metagenomes than SHAP, given that (i) *p* is usually high, and (ii) many variable interactions are expected.

We compared SHAP and endoR in regards to computational performance. The two methods were compared on RF models only, since SHAP values are computed by the xgboost R-package (46) while fitting the model instead of post-hoc. SHAP values were generated from RF using the `shap` function from the iBreakDown R-package (49). We found endoR to be substantially faster than `shap`. Specifically, endoR scales linearly with dataset dimensionality and sample size, while `shap` scales superlinearly (Supplementary Figure 8 A and C). As expected endoR CPU usage scaled linearly, increasing with the number of bootstraps (Supplementary Figure 8 E). We note that since endoR can be trivially parallelized, endoR requires less wall-time with either *B* = 10 or 25 than `shap` for the same number of threads (Supplementary Figure 8 G). endoR requires more memory than the `shap` function, but both scale sublinearly with dataset dimensionality and require only a few gigabytes for the maximum of 100 features and 2000 observations used for the evaluations (Supplementary Figure 8 B, D, and F).

In summary, based on our evaluations on all simulated datasets and phenotypes, endoR performance is comparable or better than state-of-the-art methods in regards to identifying variables and interactions of variables associated with a response variable, while generating results that are easier to interpret in shorter computation times. Altogether, endoR surpasses state-of-the-art methods for analysing metagenome data.

#### endoR rediscovers previously reported associations between cirrhosis and gut microbial composition

To illustrate the utility of endoR for microbiome studies, we applied our proposed workflow including endoR (Figure 1) to a previously published gut microbiome dataset comprising patients diagnosed with cirrhosis versus healthy individuals (41). The dataset included 130 Chinese subjects, among which 48 % were healthy, 35 % were women, with ages varying from 18 to 78 years old (mean = 45), and BMI ranging from 16 to 29 kg.m^-2^ (mean = 22). Our full model consisted of a FS step followed by the training of a RF classifier on the selected features. The model was fitted to predict the disease status (i.e., ‘healthy’ or ‘cirrhosis’), of individuals based on their age, gender, BMI, and relative abundances of gut microorganisms derived from metagenomes (Supplementary Methods). On CV sets, it had an average Cohen’s *κ* of 0.73±0.08 and accuracy of 0.87±0.04.

endoR identified 25 stable decisions that used 20 features (Supplementary Results). Many taxa used in the stable network generated by endoR were taxonomically closely related to taxa identified in the original study, with ‘closely related’ defined as the immediate parent or child taxonomic classification in the taxonomy hierarchy (e.g., the *Bacteroides* genus is the child of the *Bacteroidaceae* family, while *Bacteroidaceae* is the parent of *Bacteroides*) (Figure 5 and Supplementary Figure 9 A). Namely, *Veillonella parvula* and *Streptococcus* were confirmed as being enriched in individuals with cirrhosis (Figure 4 and Supplementary Figure 9 C and D), as observed by studies on different cohorts (12, 50). Moreover, while the *Megasphaera* genus was significantly enriched in cirrhotic individuals in the original study, endoR further identified that the species *Megasphaera micronuciformis* was the most important one to discriminate gut microbiomes of healthy individuals from those of cirrhotic individuals (Figure 4 and Supplementary Figure 9 B). The species was detected in 24 % of healthy individuals versus 85 % of individuals with cirrhosis. Additionaly, for the samples in which *Megasphaera micronuciformis* was detected, the average abundance was 10 times lower in healthy individuals compared to cirrhotic ones (respectively, 0.40±1.50·10^−4^ and 4.26±10.41·10^−4^). Intriguingly, neither the genus nor the species were identified in other cohorts (12, 50). Thus, *M. micronuciformis* may be a marker of cirrhosis specific to the Chinese cohort sampled by Qin et al. (41). We note that *M. micronuciformis* was originally isolated from a liver abscess and pus sample (51).

**Figure 5.**
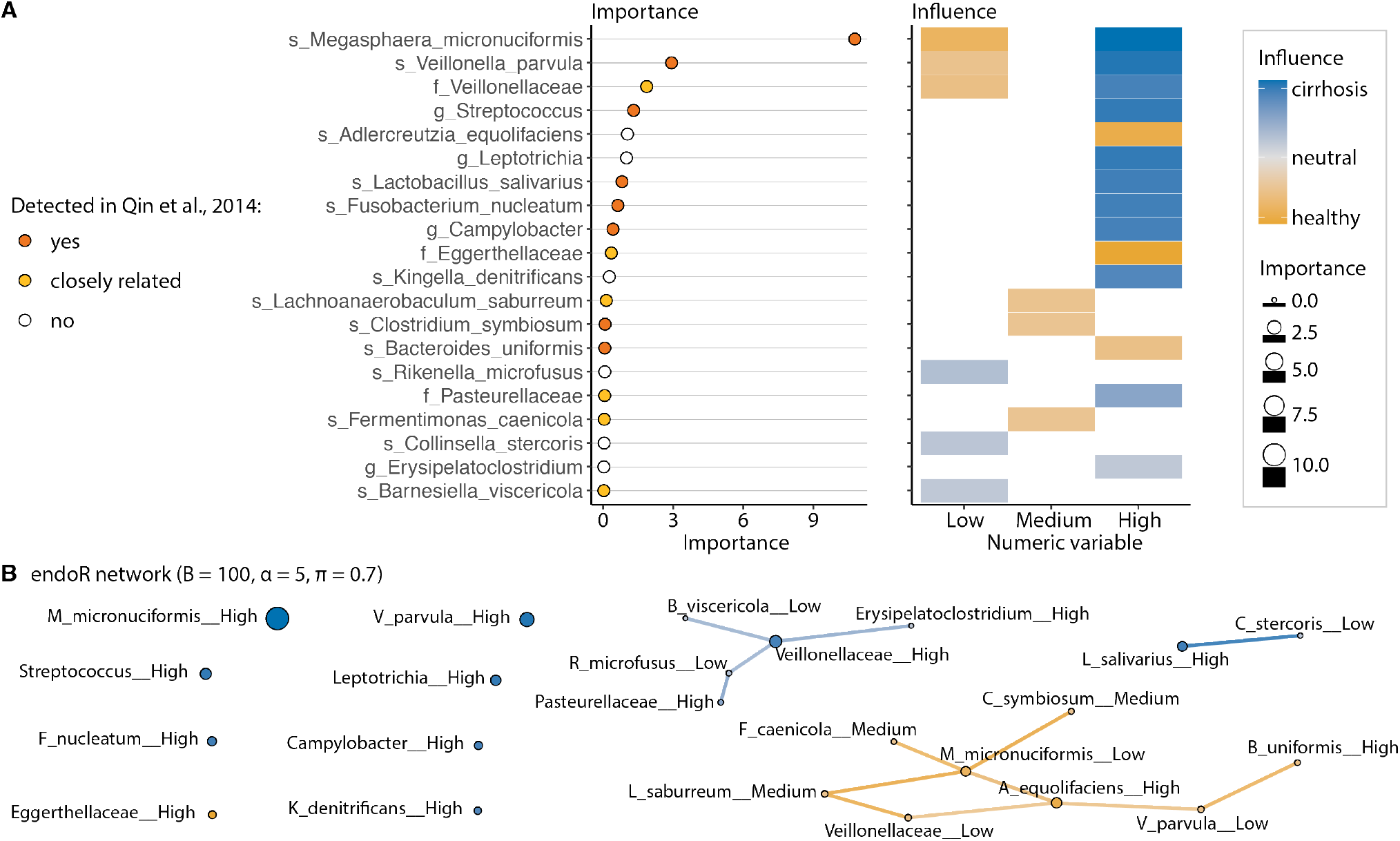
endoR recapitulates previous findings on differences in gut microbiomes between healthy individuals and patients diagnosed with cirrhosis. A/ Feature importance aggregated across each level of discretized variables and influence per-level as determined by endoR. Levels correspond to the categories a discrete variable can take, and so here to relative abundance groups created by endoR (i.e., whether samples had ‘Low’, ‘Medium’ or ‘High’ relative abundances of each taxon). ‘closely related’ designates taxa that are the direct parent or child taxonomic classification of a taxon originally associated with disease status in Qin et al. (41). White boxes in the influence plot signify that the level was not used in any stable decision; thus, the influence could not be calculated. B/ Decision network extracted from the stable decision ensembles. See Figure 2 for the description of the network; the boxed legend is shared for A and B.

Certain associations identified in the original study were not detected by endoR (Supplementary Figure 9 A). This can be partially explained by the stringent feature selection step in our model construction, which reduced the feature space from 922 to 81 taxa. For instance, significantly lower relative abundances of *Alistipes* (family *Rikenellaceae*) were found in individuals with cirrhosis by (41) and in other cohorts (12, 50). In our analysis, relative abundances of *Alistipes* were not used by the model to classify diseased and healthy samples (Figure 4), which is likely due to the large overlap in *Alistipes* relative abundance distributions between healthy and cirrhotic individuals (Supplementary Figure 9 F). However, *Rikenella microfusus*, the other genus of the *Rikenellaceae* family detected in the dataset, showed lower overlap in relative abundances between cirrhosis (depleted) and healthy individuals (enriched); thus, it was selected and used by the model (Figure 4, Supplementary Figure 9 F). In another example, the *Pasteuralleceae* family was found to be enriched in cirrhotic individuals with endoR but not in the original study (Figure 4 and Supplementary Figure 9 A and G). However, the two most abundant genera in the *Pasteurellaceae* family, the *Haemophilus* and *Aggregatibacter* genera, were identified as differently enriched in healthy versus cirrhosis individuals by Qin et al. (41) (Figure 4 and Supplementary Figure 9 A and H). In conclusion, some of the discrepancies between our analysis and the original study may be due to our use of a RF model, which can integrate nonlinear associations. Also, our feature selection step selected taxa often closely related to the genera identified in the original study, indicating that these sister taxa are actually more predictive of cirrhosis when using a RF model.

endoR identified new associations between cirrhosis and the gut microbiome. Among others, we found additional oral-microbiome associated taxa to be enriched in cirrhotic individuals. For instance, endoR revealed an important enrichment in the *Leptotrichia* genus in individuals with cirrhosis (Figure 5 A). This taxon is part of the oral microbiome (52) and is enriched in patients with periodontal disease (53). In addition, endoR identified an enrichment of the oral-taxon *Kingella denitrificans*, a member of *Neisseriaceae* (52), in individuals with cirrhosis (Figure 5 A). Altogether, these findings support the hypothesis of Qin et al. (41), in which oral commensals colonized the guts of patients with liver cirrhosis.

Our analysis also revealed an important depletion of *Adlercreutzia equolifaciens*, a bacterium associated with healthy individuals (42) (Figure 5 A and Supplementary Figure 9 E). Additionally, the decision network extracted from the stable decision ensemble contained only a few edges (Figure 5 B), meaning that a few interaction effects of bacteria on cirrhosis were found. Hence, our analysis suggests few higher order interactions among gut microbiota in relation to cirrhosis.

### New insights into the ecology of human gut *Methanobacteriaceae*

We utilized endoR to gain insight into the factors influencing the prevalence of *Methanobacteriaceae* in the human gut. We focused on this microbial clade because (i) *Methanobacteriaceae* are the most prevalent and abundant archaea in the human gut (54, 55), (ii) methanogenic archaea influence bacterial fermentation via H_2_ consumption (56–58), (iii) species of *Methanobacteriaceae* have been shown to form a complex trophic network with certain bacteria (2, 55, 58–64), and (iv) *Methanobacteriaceae* have been associated with various host phenotypes such as constipation and slow transit (65, 66), non-western diet (67–69), and body mass index (BMI) (58, 59, 70–78). Therefore, *Methanobacteriaceae* is a prime candidate for the application of endoR to resolve how this clade associates with bacterial taxa and host factors (e.g., BMI).

Metagenomes gathered for this analysis comprised 2203 individuals from 26 studies living in 23 countries across the globe (Supplementary Tables 7 and 8). Participants varied in ages from 19 to 84 years old, with a median and mean age of 33 and 40 years old, respectively. BMI ranged from 16.02 to 36.41 kg.m^-2^, with median and mean values of 23.27 and 24.03 kg.m^-2^. Women comprised 62.30 % of individuals, and 76.53 % of individuals were from westernized populations (79, 80).

We trained tree ensemble models to predict the presence of *Methanobacteriaceae* in the human gut by using taxon and metabolic pathway relative abundances, host descriptors, and metadata (see the Supplementary Tables 7 and 8, for a description of samples, host descriptors and metadata included, and Supplementary Methods, and Supplementary Figure 10, for a description of model selection and fitting). Metadata comprised the number of reads and dataset names, and both were always included for feature selection and fitting of the classifier to make sure that algorithms could correct for these variables, if necessary (e.g., in case of batch effects) (81). To evaluate the association between the presence of *Methanobacteriaceae* and host descriptors with incomplete information across samples (no age, BMI, and gender information was reported for 528, 1183, and 432 individuals, respectively), we subset observations to the 748 samples with complete information and applied our model fitting procedure. The best performing model had an average accuracy of 0.80±0.03 and Cohen’s *κ* of 0.55±0.06, based on unseen observations (Supplementary Table 9). Age, BMI, and gender were never selected across any CV set of this model. Therefore, they were excluded from further analyses and only human descriptors with complete information were included in the set of variables used to select the final model on all 2203 observations, such as country of sampling (Supplementary Table 8).

The final model accuracy and Cohen’s *κ* was 0.82±0.01 and 0.60±0.03, respectively (Supplementary Methods, and Supplementary Table 9). For the purpose of data interpretation via endoR, we trained a model on all observations and included 107 features selected by the taxa-aware gRRF algorithm as well as the metadata (Supplementary Figure 11 B).

A stable decision ensemble was extracted from the predictive model using endoR with *α* = 5 and 100 bootstrap resamples. The ensemble comprised 60 decisions that could make predictions on all samples, with an average decision error of 0.40±0.07 and support of 0.37±0.12 (Supplementary Table 10). A total of 34 features were used in decisions to predict the presence of *Methanobacteriaceae* (Figure 6 A and Supplementary Table 11). Feature importances were consistent between endoR and the mean decrease in Gini index (Supplementary Figure 11).

**Figure 6.**
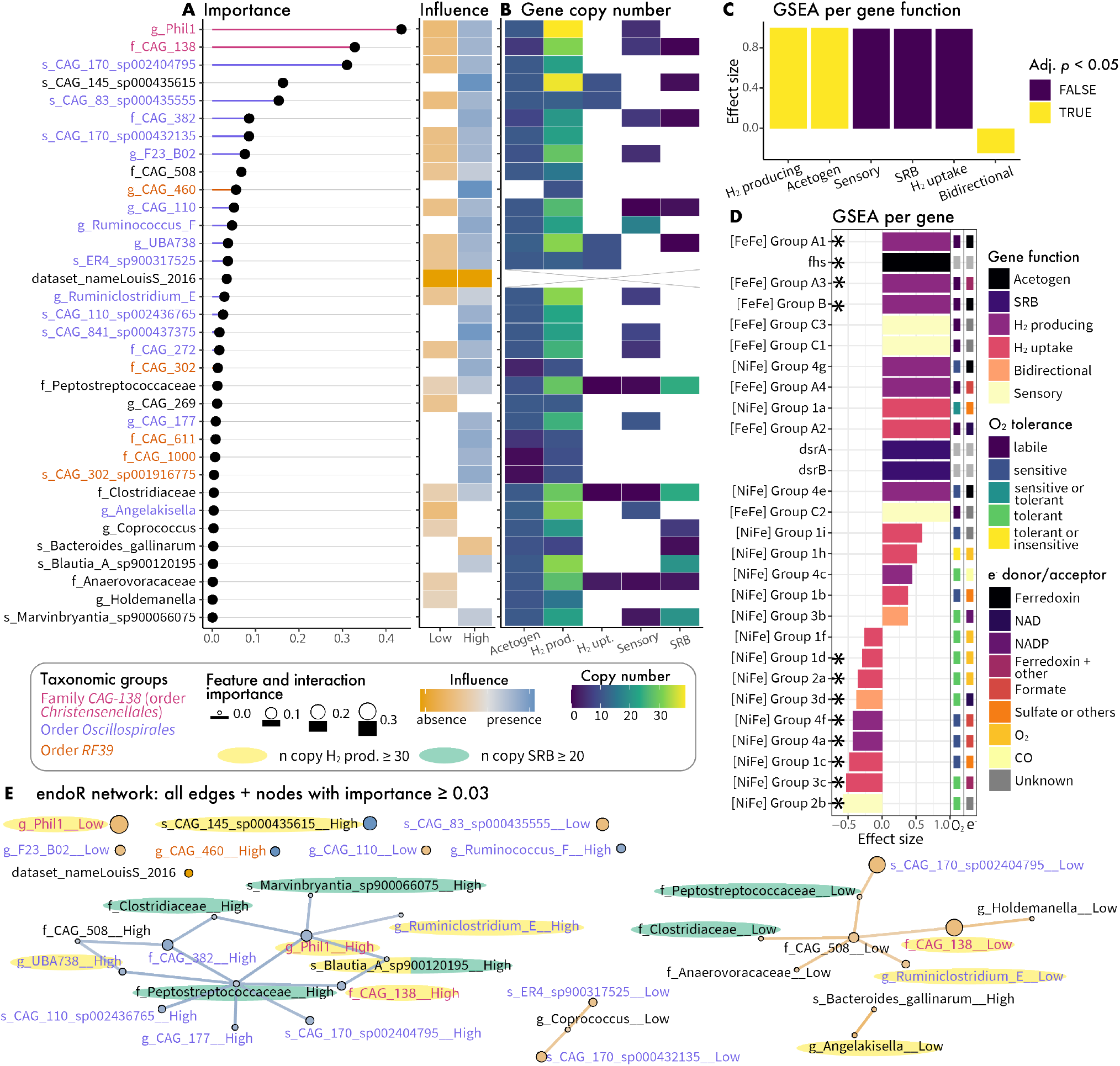
Relative abundances (RA) of *Oscillospirales, Christensenellales* and other bacteria define a gradient favorable to colonization of human guts by *Methanobacteriaceae*. A/ Feature importance and influence for each taxa used by the decision ensemble generated by endoR. Taxonomic levels are indicated with label prefixes: ‘f_’ = family, ‘g_’ = genus, and ‘s_’ = species, while taxonomic orders are indicated via bar and label colors. Levels correspond to ‘Low’ and ‘High’ relative abundances of taxa. B/ Sum of gene copy numbers of marker genes involved in H_2_ production and consumption (see Methods and Supplementary Table 13), for endoR selected features. SRB: *dsrA* and *dsrB* genes exclusively involved in sulfate reduction (82); Acetogen: *fhs* gene involved in acetogenesis (83); other categories correspond to hydrogenases predicted functions as determined by the HydDB database: H_2_ production (H_2_-prod.), H_2_ uptake (H_2_-upt.), sensory (84). Boxes are white for taxa for which genes were not detected in their genomes. The cross indicates ‘Non applicable’ (for the ‘dataset_nameLouisS_2016’ feature). C-D/ Effect sizes from gene set enrichment analyses performed at the gene function (C/) or for each gene (D/), bars are colored by the adjusted *p*-values (Adj. p). D/ Bars are colored by gene function. The predicted O_2_ tolerance of hydrogenases and electron (e^-^) donor or acceptor are indicated by colored boxes on the right of the plot (84). Asterisks denote significance (adjusted *p*-value *<* 0.05). E/ Decision network in which nodes correspond to individual features and edges correspond to pairwise interactions. Nodes and edges colors describe the feature and interaction influence; their sizes and widths are proportional to their importances. Nodes with an importance ≥ than 0.3 but not connected are shown. Taxa with a gene copy number ≥ 30 for H_2_-production and ≥ 20 for SRB genes are highlighted in yellow and green, respectively. The boxed legend applies to A, B, and E.

Both the *CAG-138* family (order *Christensenellales*, class *Clostridia*) and the *Phil-1* genus within *CAG-138* had the highest feature importances (Figure 6 A). The *Oscillospirales* order (class *Clostridia*) was over-represented in features used by endoR compared to what would be expected by random (*p*-value = 10^−3^, Supplementary Table 12), with 15 taxa from this order of the 272 taxonomic features (family, genera, and species) detected in the dataset and included in decisions (Figure 6 A). The *RF39* order (class *Bacilli*) was also over-represented (*p*-value = 10^−3^, Supplementary Table 12). Most taxa belonged to the *Clostridia* class (26 taxa, Supplementary Table 11) and had relatively higher importances compared to other features. Accordingly, the relative abundance of the *Clostridia* class was significantly associated with the presence of *Methanobacteriaceae* (Wilcoxon rank-sum test, *p*-value = 1.18·10^−20^, Supplementary Figure 12).

We note that none of the host descriptors or metabolic pathways were predictive, indicating that microbiome taxonomic composition may be more important for determining the prevalence of *Methanobacteriaceae*. However, we must acknowledge that (i) host descriptors were limited, and (ii) metabolic pathway diversity was likely undersampled. Interestingly, the model identified a cofounding effect due to possible dataset bias: samples from the LouisS_2016 study were indeed depleted in *Methanobacteriaceae*. This dataset comprised 92 stool samples from German individuals, of which *Methanobacteriaceae* was never detected. The authors utilized a non-standard DNA extraction protocol (85), which may explain the lack of *Methanobacteriaceae* detection, given that extraction protocols differ substantially in their lysis efficiency of methanogenic archaea (86, 87).

To assess whether bacterial taxa selected by endoR may be part of a H_2_-based syntrophic network, we estimated the number of genes involved in H_2_ production and consumption for the 33 taxon features (Figure 6 B). Specifically, we utilized representative genomes and assessed (i) genes coding for hydrogenases involved in H_2_ production, H_2_ consumption, both (bidirectional), or H_2_ sensing (84), (ii) genes involved exclusively in sulfate reduction (*dsrA* and *dsrB*) (82), and (iii) genes involved in acetogenesis (*fhs*) (83) (Figure 6 B and Supplementary Figure 13). To determine which of these genes were enriched among the endoR-selected features, we conducted a gene set enrichment analysis (88) based on endoR importance values. When we grouped genes by function (e.g., ‘H_2_ uptake’ or ‘SRB’), H_2_-production and acetogens were significantly enriched, while bidirectional hydrogenases were depleted (adjusted *p*-value *<* 10^−3^, Figure 6 C). In particular, 22 of the 33 taxa possessed more than 20 copies of genes coding for hydrogenases involved in H_2_-production (Figure 6 B). At the per-gene level, the acetogen marker gene (fhs), along with the H_2_-producing [FeFe] Group A1, A3, and B hydrogenases were significantly enriched, while many [NiFe] hydrogenases were significantly depleted (adjusted *p*-values *<* 10^−2^, Figure 6 E). Furthermore, there was a clear gradient of higher O_2_ sensitivity for enriched hydrogenases and increased O_2_ tolerance for depleted hydrogenases (Figure 6 D). These results suggest that *Methanobacteriaceae* co-occurs with acetogens and H_2_-producing bacteria possessing [FeFe] hydrogenases, while the negative association between bacteria possessing O_2_-tolerant [NiFe] hydrogenases suggests O_2_ exposure may be a common cause of *Methanobacteriaceae* absence.

Interestingly, the endoR decision network showed a strong positive association between *Phil-1* and the four taxa with the highest number of *dsrA* and *dsrB* gene copies: *Clostridiaceae, Peptostreptococcaceae, Blautia A sp900120195*, and *Marvinbryantia sp900066075* (Figure 6 B and E). The influence of these H_2_-consumers is not pronounced, but the interaction effect between the relative abundances of these taxa and the high relative abundances of *Phil-1* is clearly associated with the presence of *Methanobacteriaceae* (Figure 6 E). While sulfate reducers generally out-compete methanogens for H_2_ (89, 90), *Phil-1* may generate enough to alleviate H_2_ competition. Alternatively, the sulfate reducers or *Methanobacteriaceae* may be utilizing alternative substrates for growth.

## Discussion

Applying machine learning to microbiome data has increased in popularity due to the approach’s compatibility with the high-dimensional, compositional, and zero-inflated properties of amplicon and shotgun metagenome data (15, 16, 18). However, interpreting machine learning models to gain mechanistic insight into processes underpinning microbial diversity and ecosystem functioning can be challenging. We showed, through extensive validation on simulated and real microbiome data, that our proposed procedure endoR addresses these difficulties by recovering and visualizing the important components of tree based machine learning models. First, the accuracy of identifying important features and the interactions among features surpassed or at least rivaled existing state-of-the-art methods (Figure 4). Second, the feature importance and influence plots and decision networks generated by endoR were straight-forward to interpret and provided more information than existing methods (Figure 2 H-J versus Figure 2 G, Supplementary Figures 6 and 7). Third, endoR was robust to the choice of hyperparameters (Figure 3 A and D) and, by including several regularization steps (e.g., resampling inspired by stability selection (91)), effectively controlled false discoveries, even in settings with small sample sizes (Figure 3 E). Fourth, endoR is flexible: it can be applied to both random forests and gradient boosted trees, which themselves can be applied to both regression and classification tasks involving various types of features (e.g., microbial abundances and metadata). Finally, endoR is substantially more computationally efficient than, for example, SHAP (Supplementary Figure 8 A-D, and G), which is a common approach for ML model interpretation (31, 35, 92, 93).

Our re-evaluation of healthy and cirrhotic individuals initially assessed by Qin et al. (41) highlights the ability of endoR to detect known microbe-disease associations while also revealing how microbial features interact in regards to disease status (Figure 5). For example, the feature importance calculated by endoR highlighted the main microbial factors previously shown to distinguish cirrhotic and healthy individuals – particularly emphasizing the importance of *M. micronuciformis* and *V. parvula*. Notably, our approach revealed microbe-disease associations not identified in the original study. endoR found additional bacteria common in the oral microbiome to be enriched in gut microbiome of cirrhotic individuals, among which one was associated with periodontitis (53), a condition more prevalent in individuals with alcohol-related cirrhosis, presumably due to a decrease in oral hygiene (94). endoR also found *Adlercreutzia equolifaciens* to be depleted in individuals with cirrhosis (Figure 5 A). This bacterium is associated with healthy individuals compared to ones suffering from primary sclerosing cholangitis, which can lead to cirrhosis (42).

Given the importance of methanogens for mediating bacterial fermentation via syntrophic H_2_ exchange, we applied endoR to understand which bacteria and host factors determine the presence of *Methanobacteriaceae*, the dominant methanogenic clade in the human gut. Our extensive dataset, comprising a global collection of 2203 samples from 26 studies, allowed for a robust assessment across disparate human populations. endoR identified 33 bacterial clades to be predictive of *Methanobacteriaceae*’s presence. In particular, we confirmed the strong association previously observed between *Methanobacteriaceae* and members of the *Christensenellales* order (58–62), particularly with the uncultured *CAG-138* family (Figure 6 A). We also found members of the order *RF39* (class *Bacilli*) to be positively associated with *Methanobacteriaceae*. This is consistent with findings from (59) who described that *RF39* and *Methanobacteriaceae* belong to a consortium of co-occurring taxa, with *Christensenellales* forming the central hub. *RF39* are uncultivated microorganisms with very small genomes and are predicted to be acetogens (95, 96). Hence, the co-occurrence of *RF39* and *Methanobacteriaceae* may be a result of their affinity for H_2_ produced by *Christensenellales*. Nonetheless, contrary to other H_2_-consumers, no interaction effect was found between members of the *RF39* order and *Christensenellales* for predicting the presence of *Methanobacteriaceae* (Figure 6 E). As acetogenesis is a facultative metabolic pathway and *RF39* are predicted to produce H_2_ (96), H_2_ syntrophy may be an additional underlying mechanism of the association between members of the *RF39* order and *Methanobacteriaceae*.

Our findings highlight the importance of H_2_ production and consumption for predicting the presence of *Methanobacteriaceae* (Figure 6). Clades known to include acetogens and SRB were among the taxa positively associated with *Methanobacteriaceae*, which would seem to indicate competition for H_2_; nonetheless, all competitors were positively associated and seemingly can coexist (Figure 6 A-B). High rates of H_2_ production may mitigate this competition. Indeed, H_2_-producing [FeFe] hydrogenases have very high turnover rates compared to [NiFe] hydrogenases (97), and they were the only hydrogenases enriched among the endoR-selected bacteria (Figure 6 D). Moreover, the enriched [FeFe] hydrogenases are O_2_ labile (84), utilize the low redox electron carrier ferredoxin (98), and are associated with obligate anaerobes (99). This contrasts the generally O_2_-tolerant [NiFe] hydrogenases not utilizing ferredoxin and that were depleted among endoR-selected taxa (84). These findings suggest that intestinal aerobiosis may mediate the presence of both *Methanobacteriaceae* and bacteria positively associated with the clade due to the low redox required for methanogenesis, along with the O_2_ sensitivity of *Methanobacteriaceae* and the bacterial H_2_ producers possessing [FeFe] hydrogenases. The absence of both *Methanobacteriaceae* and these H_2_ producers may indicate epithelial oxygenation resulting from diseases such as IBD or ulcerative colitis (100–102). Indeed, a decline in *Methanobacteriaceae* taxa has been associated with IBD, ulcerative colitis, and Crohn’s disease (103– 105).

Intestinal transit times may also be a factor determining *Methanobacteriaceae*’s prevalence. Many of the endoR-selected bacteria are members of the *Oscillospiraceae* family whose members are predicted to have slow replication times and would thus benefit from slow transit times (106). Similarly, *Methanobacteriaceae* species have generally slow replication rates and are associated with increased transit time (107, 108). Moreover, CH_4_ can slow peristalsis (109), hence methanogenesis may be indirectly promoting the persistence of *Oscillospiraceae* species via manipulating host physiology.

Still, no host factors were predictive, including BMI, while previous work has shown associations between *Methanobacteriaceae* taxa (or methanogens assessed in aggregate) with either anorexic, lean or obese phenotypes, depending on the study (58, 59, 67–78). These contradictory findings among existing studies, and our lack of association between BMI and *Methanobacteriaceae*, suggest that population-specific or study-specific factors mediate this association. While endoR could identify such context-dependent associations, our aggregated dataset may not contain the relevant factors (e.g., diet or other lifestyle factors). Westernization status was also not predictive of *Methanobacteriaceae*, although taxa in this clade have been found to be enriched in certain non-westernized populations such as Matses huntergatherers (67, 68), traditionally agricultural Tunapuco (67), or Columbians in the midst of westernization (110). The categorization of ‘westernized’ versus ‘non-westernized’ is likely overly broad to accurately predict *Methanobacteriaceae* across disparate human populations (Supplementary Figure 14 H-K). Indeed, not all studies have shown enrichment of *Methanobacteriaceae* in ‘non-westernized’ populations (80, 111, 112).

In summary, endoR advances the state-of-the-art for interpreting machine learning models trained on microbiome amplicon and shotgun metagenome data. We note that regardless of the ML model interpretation method, poor model performance will generate misleading interpretations. Our evaluations of endoR’s accuracy in regards to the tree ensemble model accuracy provide an explicit guideline for evaluating the trustworthiness of model interpretations generated by endoR. Moreover, we provide sensible parameter defaults that will often lead to robust results, but we emphasize the careful consideration of parameters based on our extensive evaluations. As we show with our validations and application to gut-inhabiting *Methanobacteriaceae*, endoR produces robust and informative model interpretations. These allow researchers to gain insight into the biological mechanisms underpinning ML model predictive performance and help guide controlled experimentation to directly test such mechanisms.

## Methods

This section is divided into three parts: (i) a detailed description of endoR, (ii) a summary of the evaluation metrics, and (iii) an overview of the simulated and real data. Further methodological details and the technical implementation are covered in the Supplementary Methods.

### Description of endoR

endoR takes a fitted tree-based ML prediction model and extracts a regularized decision ensemble. Based on this decision ensemble it computes decision, feature, and interaction importance and influence metrics by assessing their individual contribution to the overall prediction. These metrics are visualized in easy-to-interpret plots that can be used to gain insights into the fitted model. In the following, we describe the mathematical details underlying the decision ensembles and metrics, explain how endoR regularizes the decision ensemble, and present how results are visualized.

#### Rules, decisions and decision ensembles

Let **x** = (*x*^1^, …, *x*^*p*^) ∈ ℝ^*p*^ represent *p* features (numeric or factor variables, e.g., relative abundances of taxa and host gender), *y* ∈ ℝ a response variable, and assume we have observed a sample of *n* observations (**x**_1_, *y*_1_), …, (**x**_*n*_, *y*_*n*_) ∈ ℝ^*p*+1^. Our framework is able to handle both regression (*y* continuous) and binary classification (*y* binary); endoR transforms multi-class classification tasks to binary problems (one class versus all others).

A *rule* is a function *r* : ℝ^*p*^ → {0, 1} of the form

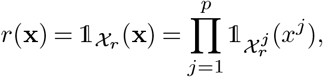

where 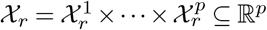. We define a *decision* to be a tuple *D* = {*r*_*D*_, *ŷ*_*D*_} consisting of a rule *r*_*D*_ and a constant *prediction ŷ*_*D*_. The prediction *ŷ*_*D*_ is computed during model fitting following any pre-defined estimation procedure (e.g., least-squares) and should be thought of as a good approximation of *y* on the *sample support S*_*D*_ := {*i* ∈{1, …, *n*}| *r*_*D*_(**x**_*i*_) = 1}, the subset of samples following the rule. Decisions are the building blocks of a large class of non-parametric ML models such as random forests and boosted trees. These models combine many decisions to construct high-capacity prediction procedures. Any such model can be seen as a collection of decisions 𝒟 = {*D*_1_, …, *D*_*M*_}, which we call a *decision ensemble*, together with an appropriate method for aggregating the predictions (20).

For every subset of observations *S* ⊆ {1, …, *n*}, we define the error function *α*(*S*, ·) : ℝ → ℝ either as the mean residual sum of squares in the case of regression, or by the mean misclassification error in the case of binary classification, formally,

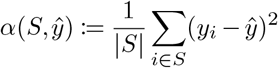

or

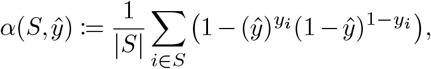

respectively. For a fixed decision *D* and a variable *x*^*j*^, or pair of variables {*x*^*j*^, *x*^*k*^}, we define the *complement decision* 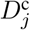, or 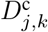, to be the decision resulting from modifying the rule *r*_*D*_ to have the complement support for the variable *x*^*j*^, or for the pair of variables {*x*^*j*^, *x*^*k*^}. Additionally, we define the decisions 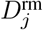 and 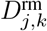 to be the decisions resulting from removing the variable *x*^*j*^, or the pair of variables {*x*^*j*^, *x*^*k*^}, from the rule *r*_*D*_. See the Supplementary Methods, and Supplementary Figure 15, for a visualization of these modified decisions. The predictions 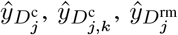 and 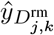 are each updated based on the new rule.

For a variable *x*^*j*^, we define the set of *active decisions* as 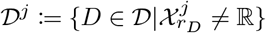, which is the subset of decisions which depend on *x*^*j*^. Likewise, the set of active decisions of a pair of variables {*x*^*j*^, *x*^*k*^} is defined as 𝒟^*j,k*^ := 𝒟^*j*^ ∩𝒟^*k*^.

#### Decision importance

For a decision *D* ∈ 𝒟, we define the *decision importance* by

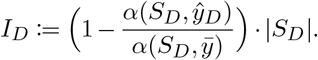

This quantifies the improvement of predicting *y* on the support *S*_*D*_ with *ŷ*_*D*_ instead of with the full sample average 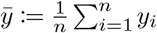. It is weighted by the size of the decision’s support.

For regression and binary classification, 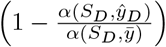 corresponds to the coefficient of determination (or R^2^) (113) and Cohen’s *κ* (114), respectively, computed on the subsample *S*_*D*_. Thus, the decision importance is a quality measure that incorporates both the support size and predictive performance of the decision.

#### Feature and interaction importance

For a variable *x*^*j*^, we define the *decision-wise feature importance*

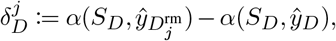

as the difference in predictive performance on *S*_*D*_ between *ŷ*_*D*_ and 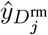 (i.e., utilizing versus not utilizing information about *x*^*j*^ for the prediction, Supplementary Figure 15).

For a pair of variables {*x*^*j*^, *x*^*k*^}, the *decision-wise interaction importance*

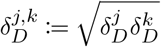

is the product of the decision-wise feature importances of *x*^*j*^ and *x*^*k*^. We use the square root to ensure that the interaction importance remains on the same scale as the feature importance.

The *feature importance* and *interaction importance*,

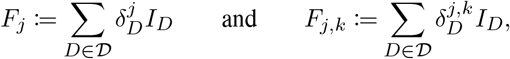

respectively, are then obtained by summing decision-wise feature and interaction importances over all decisions 𝒟 in weighted by the decision importance. High values of the feature and interaction importances indicate that the variable, or pair of variables, contribute strongly to important decisions.

#### Feature and interaction influence and direction

For every decision *D* and variable *x*^*j*^, we define the *direction indicator* 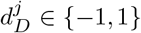 to express whether a rule predominantly uses small or large values of that variable (see Supplementary Methods). And, we calculate *η*_*j,k*_ := 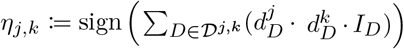 to record whether variables {*x*^*j*^, *x*^*k*^} are each associated with *y* in the same direction.

To measure the influence of a feature, or pair of features, on the prediction *ŷ*_*D*_ of a decision, we proceed similarly as with the feature importance, though we now compare actual predictions instead of errors of predictions on *S*_*D*_.

We define for a variable *x*^*j*^ and a pair of variables {*x*^*j*^, *x*^*k*^}, the *decision-wise feature influence* and *decision-wise interaction influence* as

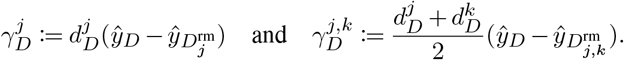

A large positive value of 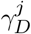 indicates that large values of *x*^*j*^ are positively associated with the response *y* on the support of the rule, while negative values of 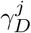 imply a negative association. Likewise, a large value of 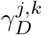 indicates that large values of both {*x*^*j*^, *x*^*k*^} are positively associated with *y*, and 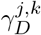 is negative when small values of both {*x*^*j*^, *x*^*k*^} are negatively associated with *y*. In addition, 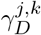 is equal to zero when the directions of association of the variables *x*^*j*^ and *x*^*k*^ with *y* are opposite.

We assess the overall *feature influence* of a feature *x*^*j*^, and *interaction influence* of pair of variables {*x*^*j*^, *x*^*k*^}, by averaging the decision-wise feature and interaction influences, respectively,

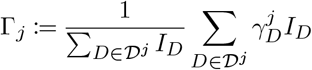

and

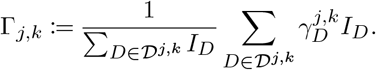

#### Regularization of the decision ensemble

We propose several procedures to regularize the decision ensemble and so reduce the noise by including a simplicity bias. Procedures are briefly introduced here but are presented in detail in the Supplementary Methods.

### Decision-wise regularization

The optional first and second steps involve discretization of numeric variables (into 2 categories by default) and pruning of rules. Pruning consists of removing variables from decisions that do not substantially participate in a decision (i.e., for which the difference in errors of the decision with and without the variable is low) (38). After each decision-wise simplification step, decisions consisting of the same rules are grouped, the multiplicity is recorded (i.e., how many decisions have been collapsed into the simplified decision) and the prediction, error, support, and importances are re-computed based on the updated rule. Lastly, the decision importance is weighted by the decision multiplicity.

### Decision ensemble stability

At this stage, the decision ensemble will often be large and still include poorly predictive decisions. In addition, the metrics (e.g., feature and interaction importances) may have a tendency to overfit. To avoid these issues, endoR implements an option to simplify the decision ensemble by running all decision-wise regularization steps and decisions metric calculations on *B* bootstrap resamples of the data (this does not include refitting the prediction model). endoR then simplifies the decision ensemble by only keeping decisions that are returned consistently across bootstrap resamples. This approach is motivated by the stability selection procedure due to (91). More specifically, for user-selected parameters *α* ∈ ℝ_*>*0_ and *π*_thr_ ∈ (0.5, 1] (*π*_thr_ = 0.7 and *α* = 1 by default), the *q* most important decisions of each bootstrap resample are recorded and those appearing in at least *π*_thr_ · *B* of the resampled decision ensembles are then selected. Motivated by the theoretical results of (91) on controlling the expected number of false discoveries (*α* corresponds to the expected number of false discoveries in this context), we select *q* to be

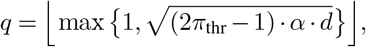

where *d* is the average number of decisions across all bootstrap resamples. For each decision in the stable decision ensemble, the decision-wise influence and importance are averaged across the resampled decision ensembles, and the influence and importance are re-computed as described above. By default, bootstrapping is performed on *B* = 10 resamples of size *n/*2.

#### Visualization: Decision network and importance/influence plot

After extacting a regularized decision emsemble and computing all metrics, endoR visualizes the results in a feature importance plot, a feature influence plot, and a decision network (summarized in Figure 1 A and exemplified in Figure 2 H-J). Both the feature importance and influence plots show only the main effects of variables that appear in the final regularized decision ensemble. For the influence plot, white blocks indicate either that the discretized level did not appear in the final decision ensemble or that it is a binary variable. In the decision network, nodes correspond to single variables and edges to interaction effects on the response between two nodes. Sizes represent feature and interaction importances, while colors describe feature and interaction influences. In addition, the edge type indicates the interaction direction, so that it is either a solid line if on average the pair of variables have the same sign (i.e., positively associated variables), or a dashed line, if not.

### Datasets

#### Simulated datasets

We generated *n* independent observations of a random vector (*Y, K, V* ^1^, …, *V* ^12^) as follows. Let *V* ^1^, … *V* ^12^ be independent 𝒩(0.5, 1) distributed random predictive variables, let *K*, a multiclass feature, be uniformly distributed over the categories {a, b, c, d}. The binary response *Y* is set by the rules in Table 1, its sign is changed with a probability *r* to add noise.

**Table 1.** Predetermined decision rules to generate the response variable from the simulated datasets.

| Decision rule | Response |
| --- | --- |
| Group = 'a' & $V^1 > 0$ & $V^2 > 0$ | 1 |
| Group = 'a' & $V^1 \leq 0$ & $V^2 \leq 0$ | 1 |
| Group = 'a' & $V^1 > 0$ & $V^2 \leq 0$ | -1 |
| Group = 'a' & $V^1 \leq 0$ & $V^2 > 0$ | -1 |
| Group = 'b' & $V^3 > 0$ | 1 |
| Group = 'b' & $V^3 \leq 0$ | -1 |
| Group = 'c' & $V^4 > 0$ & $V^5 > 0$ | 1 |
| Group = 'c' & $V^4 \leq 0$ & $V^5 \leq 0$ | -1 |
| Group = 'd' & $V^6 \leq 0$ & $V^7 > 0$ | -1 |
| Group = 'd' & $V^6 > 0$ & $V^7 \leq 0$ | 1 |

**Table 2.** Predetermined decision rules based on the making of the artificial phenotypes.

| Decision rule | Response |
| --- | --- |
| Group = 'a' & $t_1 > 0$ & $t_2 > 0$ | 1 |
| Group = 'a' & $t_1 \leq 0$ | -1 |
| Group = 'a' & $t_2 \leq 0$ | -1 |
| Group = 'b' & $t_3 > 0.1$ & $t_4 > 0.003$ | 1 |
| Group = 'b' & $t_3 \leq 0.1$ | -1 |
| Group = 'b' & $t_4 \leq 0.003$ | -1 |
| Group = 'c' & $t_5 > 0$ & $t_6 > 0$ & $t_7 > 0.01$ | 1 |
| Group = 'c' & $t_7 > 0.01$ | 1 |
| Group = 'c' & $t_5 > 0$ & $t_6 > 0$ | 1 |
| Group = 'c' & $t_5 \leq 0$ & $t_6 \leq 0$ & $t_7 \leq 0.01$ | -1 |
| Group = 'c' & $t_5 \leq 0$ & $t_7 \leq 0.01$ | -1 |
| Group = 'c' & $t_6 \leq 0$ & $t_7 \leq 0.01$ | -1 |
| Group = 'd' & $t_8 \leq 3.98 \cdot 10^{-4}$ & $t_9 > 0$ | 1 |
| Group = 'd' & $t_8 > 3.98 \cdot 10^{-4}$ | -1 |
| Group = 'd' & $t_9 \leq 0$ | -1 |

$t_i, i \in \{1, \dots, 9\}$ were randomly sampled from the variables of the metagenomes detected in more than 50% of the samples.

We used this data generating mechanism as a very simple model to evaluate our method as the underlying mechanism is fully understood here. For readers unfamiliar with abstract simulation settings, it may be helpful to think about the variables *V* ^1^, …, *V* ^12^ as (re-scaled) microbial abundances, the categories *a* to *d* as phenotypes such as age groups and the response variable *Y* as a disease indicator, with 1 and −1 encoding healthy and diseased, respectively. A single replicate of the data with parameters *n* = 1000 and *r* = 0.05 is given in Supplementary Figure 2 A-D.

When evaluating endoR on this simulated data (see Results) we used a RF model fitted with the randomForest R-package (43). Details on the fitted model and the parameter settings of endoR are provided in the Supplementary Methods.

#### Artificial phenotypes

To assess the performance of endoR under more realistic microbiome conditions, we additionally evaluated it on a real metagenomic dataset with a simulated response variable. We call this the *artificial phenotypes* data set, to stress that while the features are real metagenomic measurements, we artificially construct groups and a response variable, hence providing a known ground truth of the underlying model. The artificial phenotype designate the response variable.

We used the same collection of human gut metagenome datasets as in (115), with additional sample exclusion criteria and identical sequence processing (Supplementary Methods). The dataset comprised 2147 samples from 19 studies, with relative abundances of families, genera and species with a prevalence above 25 % (*p* = 520 taxa; Supplementary Methods).

Based on this data, we artificially constructed a multi-class phenotypic variable *K*, uniformly distributed over the categories {*a, b, c, d*} for the replicate presented in Figure 2 A-E and 8, or {*a, b, c*} otherwise. Within each group, combinations of randomly picked taxa with a prevalence higher than 50 % were used to determine the sign of the response variable *Y* (for the replicate in Figure 2 A-E and 8, see Table 2 and Figure 2 A-E; otherwise, see the pooled list of rules from which decisions were drawn in Supplementary Table 2). Noise was added by changing the group label with a probability *r*: new group labels were drawn from {*a, b, c, d, e*} for the replicate in Figure 2 A-E and 8, and from {*a, b, c, d*} otherwise.

We evaluated endoR on these artificial phenotypes based on fitted RFs and boosted trees, generated via the randomForest and xgboost R-packages, respectively (43, 46). To additionally regularize the ML models, we combined them with FS and cross-validation (CV). All details on the fitted ML models and the parameter choices for endoR are provided in the Supplementary Methods. Each model was processed with endoR using default parameters (i.e., *K* = 2, *B* = 10, and *α* = 5; Figure 3 A-C and 4, and Supplementary Figures 5 F, I-J, 3, and 4). For the replicate in Figure 2, numeric variables were discretized into 3 categories and *B* = 100 bootstrap resamples.

#### Cirrhosis metagenomes

Metadata and gut microbial taxonomic profiles from metagenomes generated by Qin et al. (41) were downloaded from the MLRepo (https://github.com/knights-lab/MLRepo, accessedon27/01/2021). The dataset consisted of 68 and 62 stools samples from cirrhotic and healthy individuals, respectively, for whom age, BMI and sex information were available (48 % healthy individuals). The formatting of metagenomes and model fitting procedure are detailed in the Supplementary Methods. In brief, rare taxa were filtered out before model fitting. FS was applied to select the most relevant relative abundances of family, genus and species taxonomic ranks; it reduced the number of taxa from 926 to 85. A full model was fitted using selected taxa and metadata (gender, age, BMI, number of sequence reads; Supplementary Methods) and was processed using endoR with default parameters, except for the discretization into 3 categories, *B* = 100 bootstrap resamples of size 3*n/*4, and *α* = 5 (Supplementary Results).

Metagenomes and associated sample metadata from a globally distributed set of studies were gathered from (40) by (115) (Supplementary Tables 7 and 8). Details on data processing are available in the Supplementary Methods. Briefly, (i) metagenome were profiled with the HUMAnN2 pipeline to obtain metabolic pathways profiles based on the MetaCyc database (116, 117) and with Kraken2 and Bracken v2.2 based on a customized Genome Taxonomy Database (GTDB), Release 89.0 created with Struo v0.1.6 (available at http://ftp.tue.mpg.de/ebio/ projects/struo/) (118–121) for the taxonomic profiles; (ii) rare taxa were filtered out and taxonomic ranks from family to species were included (*n* = 2190 taxa; 181 families, 562 genera and 1447 species; Supplementary Figure 16); (iii) relative abundances of MetaCyc metabolic pathways at the community level with complete coverage and a prevalence greater than 25 % were included (*n* = 117 pathways). FS algorithms and tree ensemble models trained to classify samples based on the presence/absence of *Methanobacteriaceae* were compared using 10 CV sets (Supplementary Methods, and Supplementary Table 9). The final model was processed with endoR with default parameters, except for *B* = 100 bootstraps and *α* = 5.

### Evaluation metrics and benchmark methods

#### Evaluation metrics

##### Simulated data

A ground truth network was extrapolated from Table 1 (Supplementary Figure 2 E). The network constructed from the final decision ensemble by endoR (Supplementary Figure 2 H) was compared to the ground truth network by counting the numbers of true postive (TP), false positive (FP) and false negative (FN) nodes and edges.

##### Artificial phenotypes

Ground truth networks were extrapolated from the procedures used to create the artificial phenotypes (for an example, Table 2 corresponds to Figure 2 G). Since the data set is made of real metagenomes, a deficit here was the lack of ground truth on associations among predictive variables, notably from the same taxonomic branch. Hence, to account for taxonomic relationships, we extended the lists of true nodes and edges to include nodes and edges from related taxa. We considered as ‘related’ taxa the direct coarser and finer taxonomic ranks, and species from the same genus. Consequently, a node identified by endoR was counted as TP if it was in the ground truth network, or related to a node in the ground truth network. If both a true node and a related taxon were identified by endoR, the TP was counted only once to prevent inflating results. The same counting was performed for edges.

##### Metrics

Based on the numbers of TPs, FPs, and FNs, standard performance metrics (accuracy, precision, recall) were calculated to evaluate networks generated by endoR. In addition, TPs and FPs were weighted by their feature or interaction importances (for nodes and edges, respectively) to calculate a weighted precision, and so estimate the magnitude of TP in the endoR results. Given a ranking over the decision importances, TP/FP curves could be constructed for nodes and edges. To do so with endoR, for a fixed *α*, we first ranked the top *q* decisions of each bootstrap according to their probability of being selected in the final stable decision ensemble (i.e., the number of occurrences across bootstraps). Networks were computed for each probability of a decision to be selected, and the probabilities of edges and nodes to be in networks were subsequently calculated. Edges and nodes were then ranked by these probabilities and TP/FP curves were constructed for endoR (Figure 3 A and D, and Supplementary Figures 3 and 5 C and F). Curves were interpolated and averaged across repetitions.

#### Comparison of endoR with state-of-the-art

The comparison of endoR against state-of-the-art methods was based on the AP simulated data and consisted of the following steps (additional details are provided in the Supplementary Methods). First, for all numeric variables in the dataset (*p* = 520 taxa), we performed a Wilcoxon-rank sum test to identify taxa enriched in samples labelled with one or the other response variable category (‘-1’ versus ‘1’), and we performed a *χ*^2^ test to assess whether group categories comprised more samples than expected from one or the other response category; *p*-values were adjusted using the Benjamini-Hochberg correction method; features were ranked by 1 − *p*-value and effect size in case of ties. Second, we divided samples according to their response variable category and used taxa relative abundances (*p* = 520 taxa) to build sub-networks for each category via the graphical lasso (48) and sparCC (47) methods, as implemented in the SpiecEasi R-package (122); features were ranked by the square of covariance matrices parameters. For each method, edges shared between the two subnetworks were filtered out. From the RF model, we also computed Gini importances (19, 25), as implemented in the randomForest R-package (43), and SHAP values (35, 123), as implemented in the iBreakDown R-package (49). We additionally trained an XGBoost model (46) on the same features selected by gRRF (*p* = 18 taxa and group dummy variables). The XGBoost model was trained with default parameters and nrounds = 10, objective = ‘binary:logistic’. SHAP values and SHAP interaction values were extracted from it using the xg-boost and SHAPforxgboost R-packages (46, 124), and it was finally processed with endoR. For Gini, SHAP and endoR, features and pairs of features were ranked by feature and interaction importances. Variables, or pairs of variables, were randomly drawn and sorted to build TP/FP curve; the process was repeated 1000 times and averaged.

### Bacterial genome analysis

Species-representative genomes from the GTDB-r89 database, which were used to obtain relative abundances of taxa in metagenomes (120), were downloaded for each species detected in the dataset before filtering. Genomes were annotated via DIAMOND blastp (125) against the following databases: (i) Fungene (82) to identify the *dsrA* and *dsrB* genes, (ii) hydDB (84) to identify genes coding for hydrogenases, and (iii) acetobase (83), to identify the *fhs* gene. The *dsrA* and *dsrB* genes encode disulforedoxins involved in sulfate-reduction and the *fhs* gene encodes the formyltetrahydrofolate synthetase involved in acetogenesis. Hydrogenases were grouped by predicted function: H_2_-production, H_2_-uptake, bidirectional, sensory (Supplementary Table 13). For each species, the number of gene copies with a percent sequence identity above 0.50 and a length coverage above 80 % was counted. For genus and family taxonomic ranks, the number of copies were averaged across species and weighted by the average relative abundance of each species in the dataset used for analysis. Patterns of gene abundances observed with absolute copy numbers were robust to differences in genome size (Supplementary Figure 13 A).

The gene set enrichment analysis was performed using the fgsea R-package (88). Taxonomic features were used as ‘genes’ and ranked by Gini or endoR feature importance, and ‘gene sets’ were defined by gene group (Acetogen, SRB, and hydrogenases predicted functions).

### Statistics

All statistical analyses were performed in R using the stats package (126). For the *Methanobacteriaceae* analysis, we measured the over-representation of taxonomic orders in the set of features used by endoR using a Monte-Carlo procedure (Supplementary Table 12). For this, we approximated the number of family, genus, and species features expected by random, given the number of features used by endoR, by randomly drawing eighteen features from the set of taxonomic features used to fit the model. We repeated the random draws 1000 times. For each draw, the number of features belonging to each order was counted. The null distribution of each order was obtained by pooling all counts across draws, and the right-tailed *p*-value of the observed count was calculated from this null distribution.

## Abbreviations

RF: random forest
FS: feature selection
CV: cross-validation
TP: true positive
TN: true negative
FP: false positive
FN: false negative
BMI: body mass index
ML: machine Learning
FSD: fully simulated dataset
AP: artificial phenotype
DNA: deoxyribonucleic acid

## Availability of data and material

All data sets, analysis and results scripts are available at https://github.com/aruaud/endoR_ data_analysis. Our method has been fully implemented as an R-package named endoR and is available for download and with its manual at GitHub (https://github.com/leylabmpi/endoR) under an MIT licence.

## Competing interests

The authors declare that they have no competing interests.

## Funding

This work was supported by the Max Planck Society.

## Author’s contributions

AR, NP, and NY conceptualized the project, analysed data, and wrote the manuscript; AR and NP designed endoR; AR and NY curated data; AR wrote the endoR R-package; RL provided funding; AR, NP, NY, and RL reviewed and edited the manuscript.

## Acknowledgements

We thank Sofia Esquivel-Elizondo for the discussions about hydrogenases and methanogens, and Jacobo de la Cuesta, Daphne Welter, and Brandon Seah for their feedback on the manuscript. We are grateful to all who make their data open and/or who contribute to open science. Open sharing of data enabled this project.

## Supplementary Methods

Here, we describe methodological and technical procedures in more details when needed compared to the main text.

### Rules, decisions and decision ensembles

For a fixed decision *D* and a variable *x*^*j*^, or pair of variables {*x*^*j*^, *x*^*k*^}, the *complement decision* 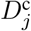, or 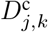, are defined to be the decisions resulting from modifying rule *r*_*D*_ to have the complement support for the variable *x*^*j*^, or the pair of variables {*x*^*j*^, *x*^*k*^} (Supplementary Figure 15), i.e.,

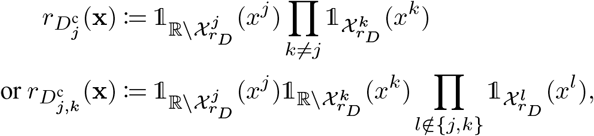

respectively.

Additionally, decisions 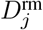 and 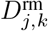 are defined to be the decisions resulting from removing the variable *x*^*j*^, or pair of variables {*x*^*j*^, *x*^*k*^}, from the rule *r*_*D*_ (Figure 15), i.e.,

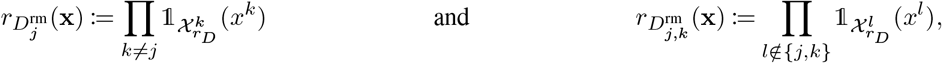

respectively.

Finally, for a subset of variables *J* ⊂ {*x*^*j*^, *j* ∈ {1,…, *p*}} the decision 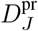 is defined to be the decision resulting from removing all variables not included in *J* from *r*_*D*_, i.e.,

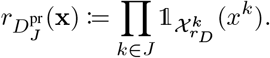

The predictions 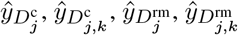 and 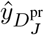 are each updated based on the new rule.

For a variable *x*^*j*^, we define the set of *active decisions* as 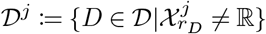, the subset of decisions which depend on *x*^*j*^. Likewise, the set of active decisions of a pair of variables {*x*^*j*^, *x*^*k*^} is defined as 𝒟^*j,k*^ := 𝒟^*j*^ ∩𝒟^*k*^.

### Extraction of rules from predictive models

Decisions are extracted from tree-based models (randomForest, ranger, gbm and xgboost (43–46)) using the inTrees R-package (38), with slight modifications. More specifically, given a tree-based model, rules are first extracted from all trees, or a subset of them, by following branches from the root down to the terminal node, e.g., for a tree composed of 4 terminal nodes, 4 decisions would be extracted.

All multi-class factor predictive variables are converted to {0, 1} encoded dummy variables. Extracted rules are then adjusted to be using only one class of each of the original multi-class factor variables and rules multiplicity is decreased accordingly. For instance, for a multi-class factor *x*^*j*^ taking values in {*a, b, c*}, three dummy variables would replace *x*^*j*^ and a rule such as “*x*^*j*^ ∈ {*a, b*}” would be transformed into two rules 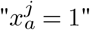 and 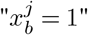 with multiplicity equal to 0.5. In addition, the same procedure of rule splitting is applied to predictive factors provided by users, that were already encoded as dummy variables for fitting the predictive model. Levels of multivariate variables are thus included only by their presence, later helping with the visualization and interpretation of networks.

### Feature and interaction direction

To understand how a single feature influences the prediction, one needs to understand whether a rule uses predominantly small or large values of that feature. For every decision *D* and variable *x*^*j*^, the *direction indicator* 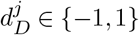

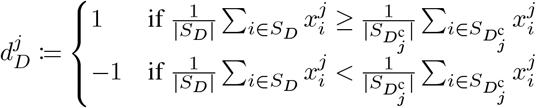

expresses whether *D* predominantly uses small or large values of variable *x*^*j*^.

For every pair of variables {*x*^*j*^, *x*^*k*^},

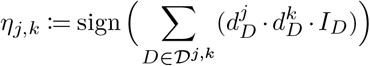

records whether variables {*x*^*j*^, *x*^*k*^} are each associated with *y* in the same direction across *D* ∈ 𝒟^*j,k*^. When both variables {*x*^*j*^, *x*^*k*^} have large, or small, values associated with the response *y*, then *η*_*j,k*_ is positive; and when large values of *x*^*j*^ are positively associated with *y* but small values are positively associated with *y*, then *η*_*j,k*_ is negative. The later occurs when 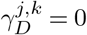.

### Regularization of the decision ensemble

We propose several procedures to regularize the decision ensemble and so reduce the noise by including a simplicity bias. These procedures are recommended but optional.

#### Decision discretization: quantiles of variable distribution

Numerical predictors can be discretized based on their quantiles (e.g., into levels ‘Low’, ‘Medium’ and ‘High’). All decisions containing discretized variables are then modified by replacing any numeric rule (e.g., ‘*x*^*j*^ ≤ *t*’) by the best approximating rule which only uses the discretized variables (e.g., ‘ *x*^*j*^ =‘Low’ ‘). Decisions consisting of the same rules are grouped, the multiplicity is recorded, i.e., how many decisions have been collapsed into the simplified decision) and the prediction, error, support and importances are re-computed based on the updated rule, and the decision importance is weighted by the decision multiplicity. Finally, the feature influence is computed for each level of discretized variables and the feature importance is calculated across all levels.

In practice, all, or a user-defined subset of, numeric variables are discretized based on their quantiles using the `discretizeVector` function from the inTrees R-package (38), adapted to accept missing values (NA). For each rule containing discretized variable, numeric thresholds are replaced by corresponding levels for which the majority of observations are included in the original sample support (Supplementary Figure 4). Rules are then transformed as described in the above section to be based on only one level, and the multiplicity is updated.

#### Decision discretization: local maxima of tree ensemble model splits

Alternatively, we propose to discretize numeric variables based on their use in the predictive model. For each numeric variable, we first collect all thresholds of splits on this variable in the model. All thresholds outside of the variable range are given the maximal or minimal values of the variable, i.e., for a threshold *t* from a split on variable *V*, if *t > max*(*V*), *t* ← *max*(*V*_*i*_) or if *t < min*(*V*), *t* ← *min*(*V*). Then, we compute their distribution and calculate the local maxima. These maxima are used as limits for the groups of the new discretized variable such that the K-1 greatest maxima are used to make K categories.

#### Decision pruning

Pruning consists of removing variables from decisions that do not participate much to a decision, i.e., for which the difference in errors of the decision with and without the variable is low (38). Comparison of errors can be performed using the absolute or relative difference in errors (absolute difference by default) (38). Accordingly, the procedure looks for the smallest subset of variables *J* with the lowest error, such as,

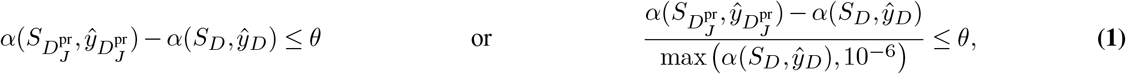

with *θ* a user-specified threshold (*θ* = 0.05 by default). If Equation Eq. (1) is not satisfied by any *J*, i.e., for all simplified decisions the differences in error are above the threshold, the original decision is returned. The prediction, error, support, importance and multiplicity are re-computed based on the updated rule, and the decision importance is weighted by the decision multiplicity.

#### Decision ensemble regularization

When bootstrapping is not feasible, we propose to instead filter out all decisions with an importance below a given threshold *λ*_imp_, selected by the user or using the following heuristic procedure

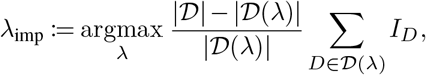

where 𝒟(*λ*) is the set of decisions with *I*_*D*_ ≥ *λ*.

### Constructing the network

After regularization and computing all metrics, we propose to visualize the feature and interaction importance and influence in a network. In particular, nodes in the network correspond to single variables and edges to interactions between variables. More specifically, for every node *j* ∈{1, …, *p*}, we choose the node size and color in the following way:

- *node size:* feature importance *F*^*j*^. Larger nodes correspond to more important variables;
- *node color:* feature influence Γ_*j*_, where the color interpolates from blue to orange (via white), with blue corresponding to small prediction values, white to prediction values close to the mean response variable across all samples, and orange to large prediction values.

Similarly, for every pair of nodes {*j, k*} ∈ {1, …, *p*}^2^, the edge between the two nodes is chosen as follows:

- *edge width:* interaction importance *F*_*j,k*_. Thicker edges correspond to more important interactions;
- *edge color:* interaction influence Γ_*j,k*_, with the same color scale than for nodes;
- *edge type:* interaction direction *η*_*j,k*_. It is either a solid line if the pair of variables is on average used in the same direction in decisions, i.e., they are positively associated, and it is a dashed line otherwise.

The network object is created using the igraph and ggraph R-packages (127, 128), hence being compatible with the broadly employed ggplot2 R-package (129).

### Implementation

We implemented the whole method described above, together with functions to visualize results, into an open source R-package available on GitHub (aruaud/endoR).

The main wrapper function of the endoR package takes as inputs (i) a predictive model fitted using the randomForest, ranger, gbm or XGBoost R-packages (43–46), and (ii) data and a response variable on which to fit the decision ensemble, being the ones used to fit the model or not. Upon starting, all factor variables are transformed into dummy variables, and, in the case of multi-class classification, the problem is transformed into a binary classification problem according to the class defined by the user to focus on. All regularization steps, i.e., discretization, pruning, filtering and bootstrapping, are optional and parameters can be elected by the user. The current implementation was optimized using the data.table R-package (130) and can be ran locally in parallel via the parallel R-package (126). Moreover, bootstrapping can be performed in parallel, locally or on a high-performance computing (HPC) environment, with the clusterMQ R-package (131).

### Data to evaluate endoR

#### Fully simulated data

Sets of simulations were performed with the following data parameters: *n* = 200, 1000 or 5000 samples and *r* = 0.05, 0.1 or 0.2 (with *n* = 1000 and *r* = 0.05 unless mentioned). Each set was replicated in 100 independent simulations (Figure 3 D-F and Supplementary Figure 5 A-E and G-H), and a single replicate of the data with parameters *n* = 1000 and *r* = 0.05 is given in Supplementary Figure 2 A-D.

We assessed the efficiency of the method to extract the right information from models by fitting an RF model on each simulation via the randomForest R-package (43) with default parameters. To tweak the accuracy of RF models for a same set of simulations and endoR parameters, we additionally the number of trees in the forest (randomForest parameter ntree = 10, 100 or 500, the default). For the replicate on Supplementary Figure 2, we report the average accuracy of models on 10-folds cross-validation (CV) 0.7−0.3 train-test. The accuracy of the model fitted on all data is reported otherwise (Figure 3 D-F, Supplementary Figure 5 A-E and G-H, and Supplementary Table 1).

Each classifier was processed with endoR using default parameters, *B* = 100 bootstrap resamples with *α* = 5 for the replicate in Supplementary Figure 2, and *B* = 10 with *α* = 5 for the replicates in (Figure 3 D-F and Supplementary Figure 5 A-E and G-H).

#### Artificial phenotypes

Data consisted of a subset of the metagenomes used in Youngblut et al. (115), so that samples with the following reported information were removed: i) samples from rectal swabs; ii) samples from individuals suffering from mumps, coeliac disease, gestational diabetes, cholera or with high relative abundances of *Vibrio cholerae*, infected by shiga toxin-producing *Escherichia coli* or cytomegalovirus; iii) samples with less than a million of sequence reads; iv) samples with missing age information. In total, metagenomes from 2147 samples from 19 studies and 23 countries were gathered. Microbial relative abundances were generated by Youngblut et al. (115) using a custom database based on the Genome Taxonomy Database (GTDB), Release 89.0 (120), created with the Struo pipeline (121). Only the relative abundances of families, genera and species with a prevalence above 25 % were included (*p* = 520 taxa).

We fitted a model predicting each artificial phenotype with the relative abundances of microbial families, genera and species, along with the multi-class group factor *K*. Note that multiple taxonomic levels were included to mimic a situation where no prior knowledge on associations between the microbiome composition and phenotype is available. Model fitting consisted of a feature selection step followed by the fitting of the random forest classifier. Feature selection was included in CV to reduce colinearity among predictors, noise and *in fine*, the dimensionality of data. Methods for models fitting are detailed below. The average Cohen’s *κ* of models from 10-folds CV 0.7-0.3 train-test sets are reported in Supplementary Table 3.

Each model was processed with endoR using default parameters and *α* = 10. For the main replicate, we used discretization into 3 categories and *B* = 100. For the repetitions on phenotype simulations, we varied *B* to be 10, 50 or 100 and *α* to be 1, 5, 10, 15 or 20 (Figure 3 A and Supplementary Figure 5 F).

#### Cirrhosis metagenomes

Metadata and gut microbial taxonomic profiles from metagenomes generated by Qin et al. (41) were downloaded from the MLRepo (https://github.com/knights-lab/MLRepo, accessed on 27/01/2021). The data set consisted of 130 stools samples from cirrhotic and healthy individuals, from which gDNA was extracted and sequenced via an Illumina HiSeq sequencer. The metagenomes had been taxonomically profiled with BURST (132) and Prokaryotic RefSeq Genomes. The downloaded taxonomic profiles consisted of read counts for taxonomic levels not collapsed at coarser level (i.e., if a read count had been assigned to the species level, the number of count of the genus was not indicated). Consequently, we calculated the read counts for each taxonomic level by summing read counts of all species in the clade. Relative abundances were then normalized per the total number of reads to obtain relative abundances. Taxa were filtered as described below.

### Progressive filtering of rare taxa

Taxa with low mean abundance or low prevalence were filtered out if, for a taxa *t* of prevalence *P* and average abundance *A*:

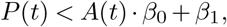

with:

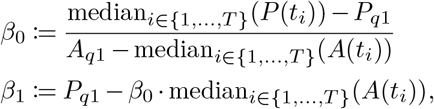

and, *P*_*q*1_ and *A*_*q*1_ corresponding to the prevalence and abundance quantile values for 25% of all taxa. This continuous filtering allows to keep taxa with low abundances but high prevalence, and inversely keep highly abundant taxa present only in a small set of samples.

### Feature selection and fitting of models on metagenome data

Microorganisms are named according to a taxonomy that divides them into groups arbitrarily defined and not consistently reflecting metabolic capacities or specificities (22, 133). Consequently, describing microbial diversity with a unique taxonomic level may not capture microbial interactions in a community. Accordingly, here we included relative abundances of the family, genus, and species taxonomic ranks. The subsequent drawback is the high-dimensionality of the data, i.e., the high number of predictive variables *p* relative to the number of observations

*n*. As *p* increases, the set of *n* observations will represent a relatively smaller set of the *p*-dimensions space (20). It will thus be harder for models to evaluate the general association of each variable or even interactions of variables with the phenotype, and associations detected may be true only for the specific set of samples used for analyses (i.e., the model will overfit data). While RF are more robust to overfitting thanks to the high number of trees built on sample bootstraps, they are more sensitive to the input feature due to the growth of trees on subsets of variables (19, 20). If many irrelevant features are included, the probability to select the true predictive ones will decrease.

Therefore, feature selection (FS) was performed before fitting an RF model to select the most relevant variables and hence improve model training. The RF was fitted with default parameter (43). A boosted tree model was alternatively fitted instead of the RF using the XGBoost R-package (default parameters and nrounds = 10) (46). The choice of the FS algorithm and parameters was determined using 10 CV with a 0.7−0.3 train-test split of the data: the model that resulted in the highest average Cohen’s *κ* was selected and a final full-model was then refitted to the entire data (Table 3).

**Table 3.**
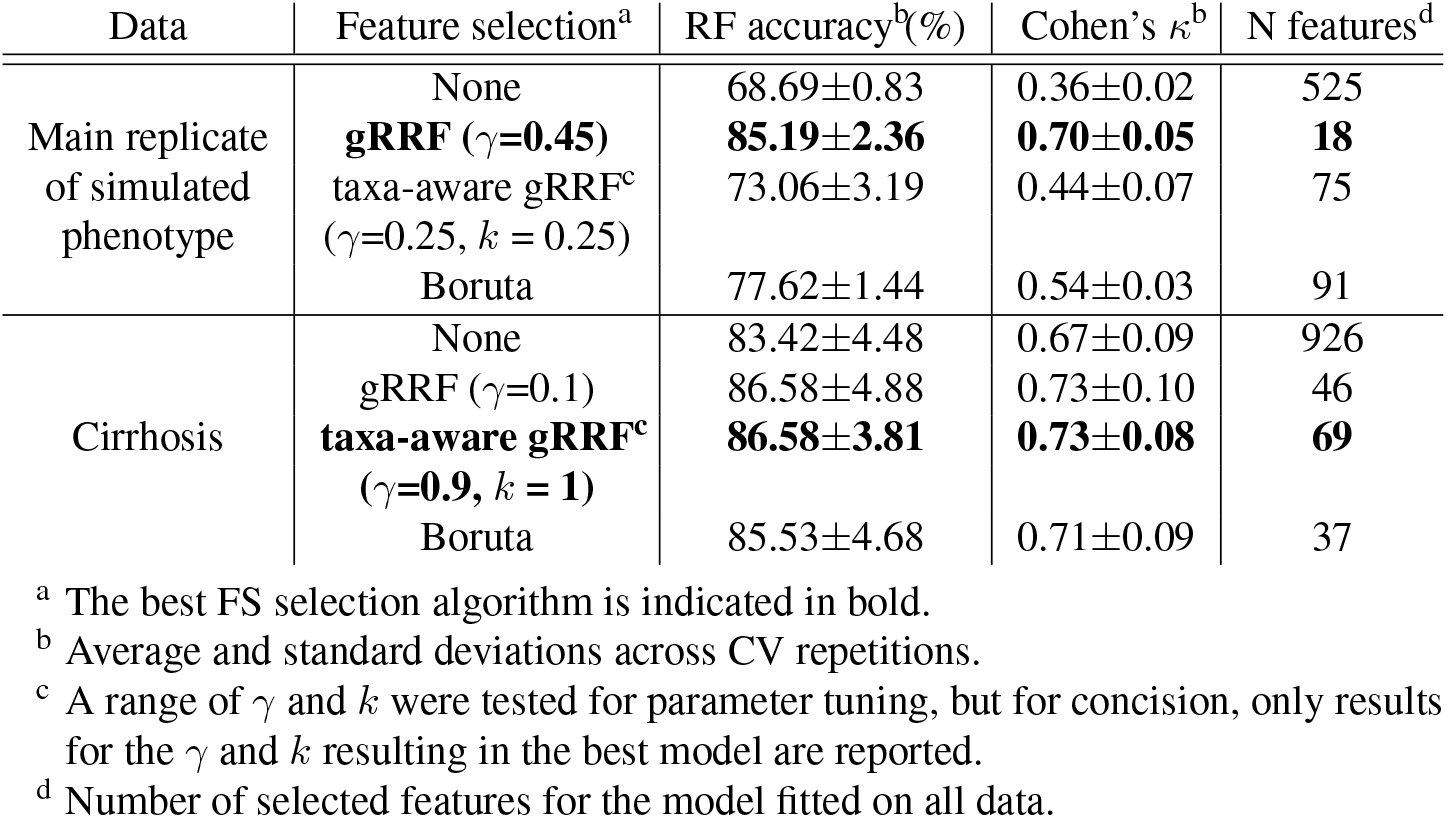
Cross-validation (CV) of feature selection and training of classifiers on metagenomic data.

The types of models considered for the metagenome experiments were the following:

- `randomForest` function from the randomForest R-package (no FS);
- subselect variables using the Boruta R-package (both functions `Boruta` and `TentativeRoughFix` with default parameters) and then apply `randomForest` from the randomForest R-package;
- subselect variables using the gRRF algorithm from the gRRF R-package for values of *γ* between 0 and 1 and, for each set of features selected with a different *γ* value, apply `randomForest` from the randomForest R-package;
- subselect variables using a modified version of the gRRF algorithm to take into account the taxonomy (see the following section), for values of *γ* and of *k* between 0 and 1 and, for each set of features selected with a different (*γ, k*) couple, apply `randomForest` from the randomForest R-package.

The choice of the Boruta and gRRF algorithms was motivated by the ability of Boruta to select all relevant variables (24), hence most likely to include all correlated variables, and for the ability of gRRF to select only relevant and non-redundant variables (21). We additionally modified the expression of the regularization term in the gRRF algorithm, to account for the hierarchical taxonomic structure in metagenomes (in the following *Taxa-aware feature selection* section, Supplementary Methods).

### Taxa-aware feature selection

Due to the hierarchical structure of nested taxonomic levels, and so their inter-dependency, redundancy occurs when including several taxonomic levels in analyses (15). But, as prior knowledge on which taxonomic levels are the most relevant is limited, it can be delicate to choose which ones to include. Nonetheless, the noise added by the inclusion of several taxonomic levels can be removed by taking the hierarchy of features into account when performing feature selection (22, 134, 135). Here we propose to modify the gRRF feature selection algorithm (21) to consider the taxonomic structure. For this, we simply add a term reflecting the importance of taxa taxonomically related to the focal one *i* when calculating its regularization term *λ*_*i*_. Hence, the original *λ*_*i*_,

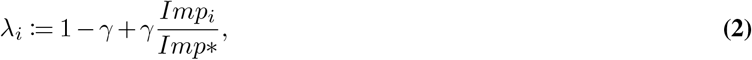

becomes

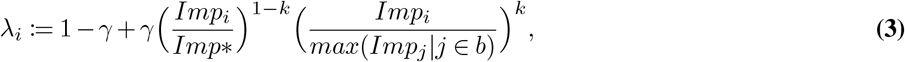

with *b* the subset of variables in the same taxonomic branch than variable *i*. For variables not describing a taxon, e.g., a metadata, *λ*_*i*_ remains calculated according to Equation Eq. (2) and not Eq. (3).

In the present article, we define *b*, as all taxa directly up- and downstream the focal one. To the finest taxonomic level used in analyses, we add the sister levels to *b*. Therefore, since we here included the family, genus and species levels to our analyses, *b* was defined for each level as:

- family: the family and all its genera;
- genus: the genus, the family it belongs to and all its species;
- species: the genus it belongs to and all species of that genus.

Both *γ* and *k* were tuned to evaluate how much weight should be given to gRRF Gini importances and to the taxonomic term in Equation Eq. (3), respectively. For each model sequence, 121 combinations of FS parameters were tested. For instance, although including the term improved models accuracy for the prediction of cirrhosis from metagenomes, it did not for the prediction of our simulated target (Table 3).

### Comparison of endoR with other analysis methods

#### Accuracy in identifying relevant variables and interactions of variables

We compared the performance of endoR in identifying true variables and interactions of variables with those of other methods commonly used for metagenome analysis.

The statistical tests used to correlate biological variables to a phenotype often are the Wilcoxon-rank sum and *χ*^2^ tests, for numeric and categorical variables respectively. Those non-parametric tests are preferred over parametric ones due to their looser assumptions on data (e.g., variable distribution). For all numeric variables in the metagenomes (*p* = 520 taxa) we performed a Wilcoxon-rank sum test to compare taxa relative abundances in target categories against each other, and we performed a *χ*^2^ test to assess whether groups were counting more samples than expected from one or the other target category. All *p*-values were adjusted with the Benjamini-Hochberg correction method. To assess the performance of non-parametric tests, variables were ordered by increasing adjusted *p*-values and were sequentially added to calculate the number of TP and FP variables.

Sparse covariance matrices are used in microbiome science to determine conditionally non-independent taxa and build correlation networks (48). The comparison of networks computed for distinct sample groups allows one to identify different associations in different groups of samples. For instance, by comparing networks extrapolated for samples collected from environment A versus environment B, one can to infer associations of variables specific to each habitat (47). A drawback of covariance matrices is the exclusion of categorical variables from analysis. Hence, for our application to the simulation on metagenomes, we could not include the group variable to the analysis. We computed the covariance matrices of samples within each target category and selected all edges not shared between the two sub-networks to estimate the accuracy of the identified edges. For each pair of variables, the square of the matrix parameter was calculated (to obtain the magnitude of the correlation between variables), pairs of variables were ordered in descending values of the square parameter, and sequentially added to calculate the number of TP and FP. Several methods exist to estimate covariance matrices, with all proposing different approaches to deal with the compositionality and expected sparsity in metagenome data. Here, methods implemented in the SpiecEasi R-package (122) were employed:

- the sparCC algorithm, which calculates the variance among observations of differences in log-transformed pseudocounts of taxa relative abundances, to estimate the covariance matrix (47);
- the (136) algorithm, which fits Lasso models on each pair of variables and uses the estimated penalization parameters to make the covariance matrix;
- the graphical Lasso method (48), which similarly to (136) fits Lasso models and uses the estimated penalization parameters to make the covariance matrix but repeats models fitting such as to maximize the log-likelihood of variables to follow a Gaussian distribution.

Due to the low accuracy of the (136) method on our simulation on metagenomes, results are not discussed nor shown in the present paper. This was expected as the (136) method is a simpler approximation of the covariance matrix, as suggested by (48).

Finally, we compared endoR to methods for interpretation of tree-based models. The most straightforward approach was to use the Gini importance (19, 25) from the same RF classifier we used for endoR, as implemented in the randomForest R-package (25). Then, we computed SHAP values (35, 123), as implemented in the iBreakDown R-package (49), with the default number of 25 random paths. SHAP estimations were then averaged across random paths for each sample and variable. For each variable, its absolute values across samples were finally averaged to obtain global SHAP values.

Since implementations of SHAP for RF classifiers in R do not return interaction values (see (49, 137) and the fastshap R-package), we additionally fitted an XGBoost model (46) to the metagenomes and artificial phenotype, with default parameters and *nrounds* = 10. SHAP values and SHAP interaction values were directly extracted from the XGBoost model by setting the `predcontrib` parameter to TRUE when fitting the model. Figures of SHAP values were created using the SHAPforxgboost R-package (124) (Supplementary Figure 7). The fitted XGBoost model was also processed with endoR for comparison.

For both SHAP and Gini importance methods, variables were sorted in descending order of feature importance (respectively the SHAP value and Gini importance) and sequentially added to calculate the number of TP and FP.

#### Computation time

We measured the computation time and memory needed to run endoR on a single replicate of the artificial phenotype data; results were compared to the `shap` function from the iBreakDown R-package (49) for comparison (Supplementary Figure 8, and Supplementary Table 4). We focused on RF for comparison with SHAP, as SHAP can be directly extracted from an XGBoost model (46), hence not requiring any additional processing time. Runs were performed in triplicates for the measurement of the total CPU time and maximal virtual memory used at any time, with 5 replicates for the wall-time. The same RF model as in Figure 2 was processed with endoR and `shap` using different sample sizes, *n* = 500, 1000 or 2000, and number of bootstraps for endoR, *B* = 1, 10, 20, 40 (Supplementary Figure 8 C-F). Furthermore, we increased the number of variables used in the predictive model by including non-selected features and fitting a new RF model via the randomForest R-package with default parameters (43) (Supplementary Figure 8 A-B). By default, we set *n* = 1000 and *B* = 1 bootstrap of size *n/*2 for processing with endoR, and SHAP values were calculated with the default parameters of the `shap` function (49). Finally, the original model with 18 features and *n* = 2147 samples was processed with endoR and `shap`, with parallelization of calculations across 4 or 10 workers (controlled by the parallel R-package). For endoR, booststraps were also allowed to be run individually in parallel using the clustermq R-package (131) (option clustermq.scheduler = “multiprocessor”). Wall-times were measured from runs on a machine equipped with Intel(R) Xeon(R) E5-4620 v4 @ 2.10GHz CPUs (80 CPUs in total).

### Investigation of *Methanobacteriaceae* in human gut metagenomes

#### curatedMetagenome database

Data used in this chapter were downloaded from the curatedMetagenomic database (40), and all samples from Youngblut et al. (115) were included, except for samples meeting the following additional exclusion criteria: (i) samples from rectal swabs; (ii) from individuals older than 90 years old, with a BMI greater than 40 kg.m^-2^, with any reported disease, or not part of control cohorts; (iii) samples from David et al., 2015 (138), due to the infection of all individuals with *Vibrio cholerae* or enterotoxigenic *Escherischia coli*; (iv) samples with less than a million sequence reads.

Information about sampled individuals comprised: country of origin, age, BMI, and whether the individual was from a westernized population. Here, westernization should be understood as an urban lifestyle with a diet composed of fewer carbohydrates and enriched in fat, sugar, and animal products compared to rural populations (79, 80). The dataset consisted of 2203 samples from 26 studies and 23 countries, among which 748 samples had complete gender, age, and BMI information (Supplementary Tables 7 and 8). We additionally grouped samples based on regional geographic origins, e.g. African countries grouped into Africa (Supplementary Table 7).

#### Enterotype clustering

Enterotypes were determined as described in Arumugam et al. (64): the Jensen–Shannon distance matrix was calculated from the relative abundances of genera using the ape (139) and phytools (140) R-packages, and partitioning around medoid was then performed with the cluster R-package (141).

#### Metabolic pathways formatting and filtering

Relative abundances of metabolic pathways were downloaded from the curated-Metagenomic database (40), where they had been obtained via the HUMAnN2 pipeline (6). All engineered, unmapped and unintegrated pathways were removed. Furthermore, only relative abundances of pathways at the community level, i.e., calculated from all gene abundances in the sample, were considered for analysis. Accordingly, we removed all relative abundances calculated from species-level gene abundances, i.e., the abundances attributed to distinct species (6). We additionally converted pathway abundances to 0 if their coverage was equal to 0. The HUMAnN2 pipeline calculates a confidence score that indicates whether reactions of pathways with non-zero relative abundances are confidently detected. A pathway coverage of 0 means that although genes coding for proteins involved in this pathway were detected, not all reactions of the pathway were confidently mapped (6). For this reason, for each sample and metabolic pathway, the relative abundance was replaced for 0 if the coverage was null. Finally, all pathways present in less than 25 % of samples were removed. A total of 117 metabolic pathways were included in analysis.

#### Taxa abundances filtering

We performed multiple taxonomic filtering steps to reduce sparsity, taxonomic redundancy, and ultimately the number of variables in the dataset.

### Filtering of rare taxa

We applied the same progressive filtering as aforementioned, on the pooled family, genus, and species taxonomic ranks. It allowed us to reduce the set of taxa from 3444 to 2318 (Supplementary Figure 16).

### Filtering of correlated taxa from a same taxonomic branch

To limit redundancy in relative abundances from taxonomic ranks of a same branch, we filtered out taxa that were significantly correlated to their direct coarser level (22). A Spearman test was performed between the two taxa, and the finer one was removed if *p*-value *<* 0.05 and *ρ*^2^ ≥ 0.95. A total of 89 taxa were filtered out in this manner.

#### *General workflow: prediction of the presence of* Methanobacteriaceae

We looked for associations between taxa and metabolic pathways relative abundances, and metadata, with methanogen presence. Their occurrence was defined as a non-zero relative abundance of *Methanobacteriaceae*.

We fitted random forest models via the ranger R-package (44), using the case.weights parameter to account for data imbalance and with the number of trees varying in {250, 500} trees. Gradient boosted models were fitted via the XGBoost R-package (46), with the number of rounds varying in {10, 50, 100, 250, 500, 750, 1000, 1500} and the maximal depth in {1, …, 10}.

The performance of models were evaluated via 10 cross-validation (CV) with 70-30 % train-test splits. Model processes were fitted to training sets and predictive performance was measured using Cohen’s *κ* on test sets (Supplementary Table 9).

For model selection, we restrained model complexity by taking into account the number of features used for fitting models and the number of trees in the forest. Decreasing the number of trees in the forest only negligibly diminished models’ performance, the best model with *ntrees* = 500 had a Cohen’s *κ* of 0.6024±0.0205 while the one with *ntrees* = 250 had a Cohen’s *κ* of 0.6004±0.0223. The best model with *ntrees* = 250 was on average using 332.50±7.79 selected features. All next seven models were also using more than 270 features on average. However, the ninth best model in term of Cohen’s *κ* used only 123.9±4.77 features and had a Cohen’s *κ* very similar to the best model (Cohen’s *κ* = 0.5957±0.0253).

#### Sets of predictors

We fitted models on different sets of predictors to reduce dimensionality. Since gender, age, and BMI were incomplete (Supplementary Table 8), we first assessed whether those variables were selected and used in models fitted on the 748 samples with complete information (*n*_*T*_ = 500). Otherwise, models were fitted on all samples without gender, age, and BMI. The metadata used as predictors were thus reduced to: country, region (i.e., the countries grouped by world region), westernization, enterotype (Supplementary Table 8). The original dataset name and the number of reads of each sample were included to each model processing step, even if they were not selected during FS.

To reduce noise and dimensionality, the taxa-aware FS step described above was performed prior to fitting predictive models.

#### Model interpretation with endoR

The final fitted model was processed with endoR: variables were discretized in *K* = 2 categories, bootstrapping was performed on *B* = 100 resamples, and *α* = 5.

## Supplementary Results

### Evaluation of endoR

#### Higher number of boostraps reduce overfitting

We generated 100 FSDs and 50 APs datasets and processed them with endoR with *B* = 10, 50 or 100 for FSDs (Figure 3 D) and with *B* = 10 or *B* = 100 for APs (Supplementary Figure 5 F). Varying the number of bootstrap resamples between 10 and 90 did not affect the precision and recall of endoR (Supplementary Figure 5 F), although higher bootstrap numbers decreased the overfitting of results (Supplementary Figure 3). This consistency in results suggests that (i) on average, endoR results are similar for different number of bootstraps, and (ii) our stability selection procedure is efficient at discriminating relevant decisions. However, increasing the number of bootstraps aids with obtaining steady decision ensembles. This slight decrease in variance, given higher number of bootstraps, is exemplified in a subsequent analysis, where we repeatedly processed replicates of the artificial phenotypes using distinct bootstrap resamples, for *B* = 10 or 100 bootstraps each time (Supplementary Figure 3). Therefore, although endoR outputs similar results regardless of the number of bootstraps, those results are more likely to be closer to the expected average results with higher number of bootstraps.

#### Discretization only marginally affects endoR

Finally, we assessed the effect of discretization. This optional step eases the interpretation of endoR outputs by simplifying continuous variables into categorical ones (e.g., ‘Low’ versus ‘High’ values grouped together). However, as this operation results in numeric variables being replaced by categories in decisions, the support and prediction of decisions are also affected. Consequently, all downstream endoR regularization steps may generate alternative stable decision ensembles for different discretization procedures. We processed the 50 APs and 100 FSDs with endoR using a discretization in *K* = 2 or 3 categories based on the distribution of each numeric variable. Our simulations showed that the precision of endoR results were similar (Supplementary Figure 5 H and J). Nonetheless, the recall was slightly higher when discretization was performed in three categories (Supplementary Figure 5 G and I), likely due to the thinner mapping of categorical variables to their original numeric ones with more categories. Increasing the number of categories also means that higher computation resources are needed: each decision may be multiplied by a factor 1 for *K* = 2 but by a factor up to 2^*pD*^ for *K* = 3, *p*_*D*_ the number of variables in the decision *D*. We note that several methods exist to discretize data. While here, and in the rest of the article, we used the quantiles of variables’ original distribution to discretize data, we propose an alternative method in the Supplementary Methods, that we compare to the present one in the Supplementary Results, and Supplementary Figure 5 E-J.

Alternatively, numeric variables can be discretized using the input predictive model. For each variable, the thresholds used in the model to make splits are employed to define the new variable categories (see Supplementary Methods). For neither of the FSD and AP datasets did the method affect endoR results (Supplementary Figure 5 E-J). Both methods are available in our package under the parameter ‘mode’, with discretization based on data distribution as default.

### New insights into the ecology of human gut methanogens

#### Individuals weakly cluster into enterotypes

We explored the spread of samples along the enterotype landscape (2, 64). Similar to previous findings (2, 64), the Jensen–Shannon distance calculated from the relative abundances of genera separated observations according to gradients of enrichment in *Bacteroides* and *Prevotella* (Supplementary Figure 14 A-B). However, samples did not strongly cluster, as shown by the within-group silhouette scores below 0.5, indicating weak clustering (2, 142) (Supplementary Figure 14 D-G). This was to be expected due to the heterogeneity of studies included in the meta-analysis and is consistent with the low silhouette scores reported for these same data (40). Clustering in three groups resulted in sample groups consistent with the ETB, ETF, and ETP enterotypes previously reported as mapping onto the gradients in *Bacteroides* and *Prevotella* relative abundances (2, 64) (Supplementary Figure 14 A-C). Since the ETF enterotype has been positively associated with higher relative abundances of *M. smithii* (2), despite the enterotypes low homogeneity, they were included in further analysis to verify their association with the methanogen.

## Supplementary Figures

**Supplementary Figure 1.**
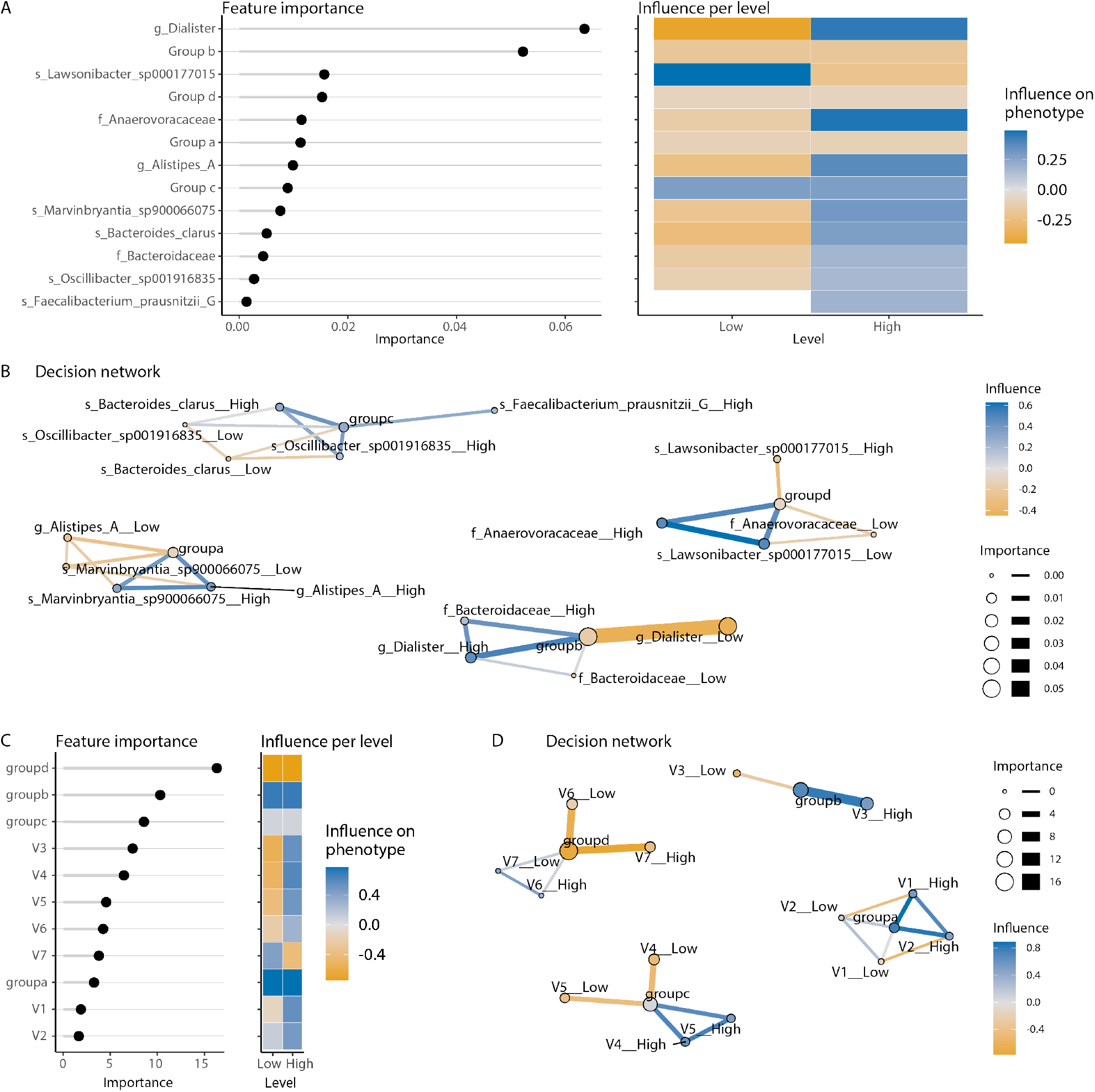
endoR recovers ground truth network from perfect predictive models. endoR was applied to the set of rules directly obtained from the true mechanism generating the response variables for one replicate of each of the AP (A-B) and FSD (C-D) simulations. No regularization step was performed, i.e., no pruning nor bootstrapping. Respective ground truth networks are visualised in Figure 2 F and Supplementary Figure 2 F. The additional edges on B are due to the discretization step. No additional edge appears on D due to the proximity between the median of numeric features (used to discretize data) and the thresholds used to make the response variable.

**Supplementary Figure 2.**
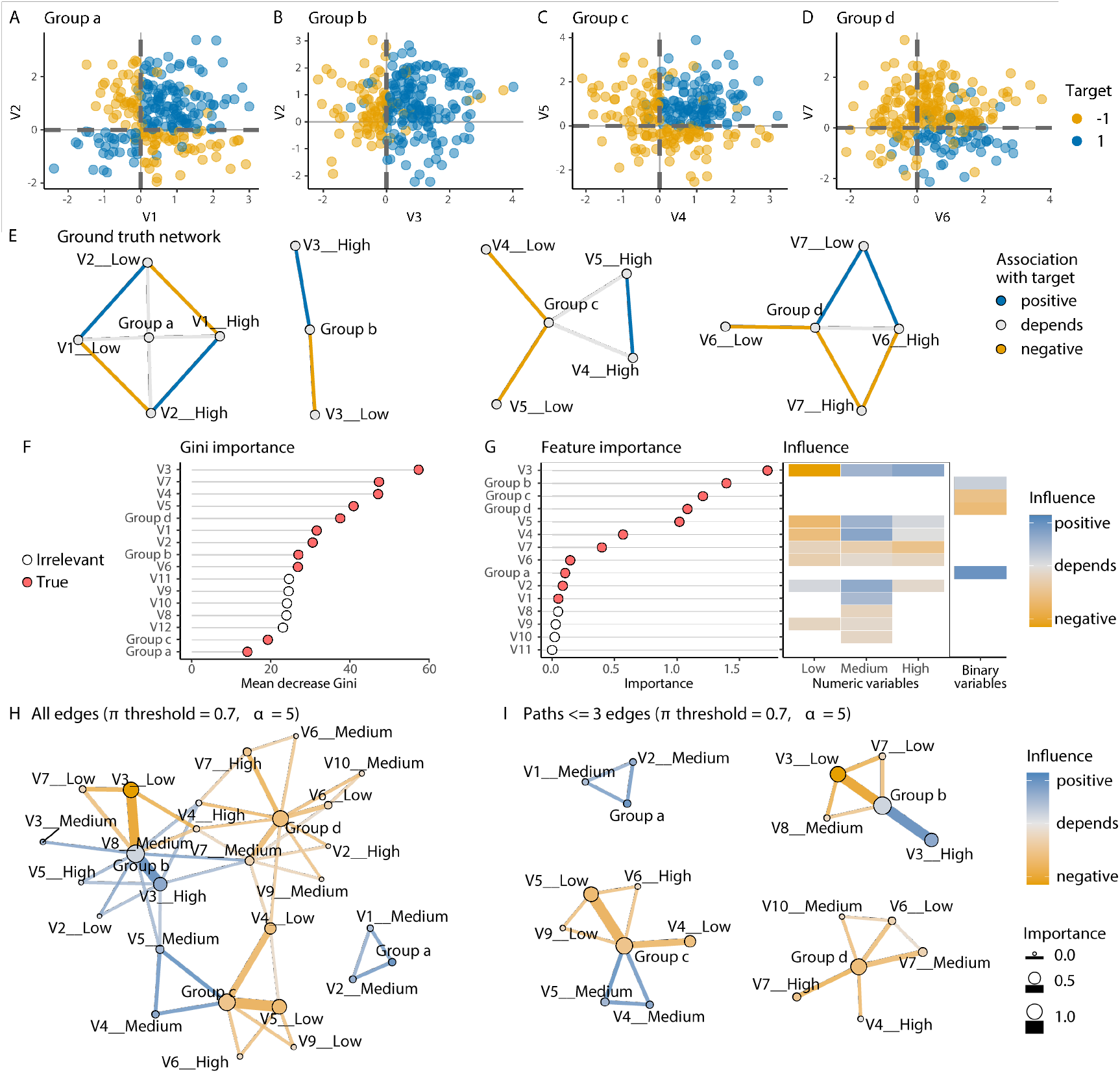
endoR captures interactions predictive of a response variable from a random forest fitted on simulated data. A-D/ Fully simulated data (FSD) structure: four groups of samples (labelled from a to d) were generated so that for each group, the binary response variable takes the value ‘1’ (blue) or ‘-1’ (yellow) according to a combination of variables described in Table 1 (e.g., V1 and V2 for Group a). The values of the response variable were then randomized with a probability *r* = 0.05. E/ Ground truth network of associations between the response variable and single variables (nodes) and pairs of variables (edges) described in A/ (see Methods). Pairs of variables predicting ‘1’ are linked by a blue edge (‘positive’) and those predicting ‘-1’ by a yellow edge (‘negative’). Variables for which high values are predictive of ‘1’ have a blue node color (‘positive’) and a yellow node color if high values are predictive of ‘-1’ (‘negative’). If high values are predictive of ‘1’ or ‘-1’ depending of other variable values (e.g., Group b predicts ‘1’ if V3 takes high values, but ‘-1’ if V3 has low values), the color is grey (‘depends’). F/ Feature importance as measured by the mean decrease in Gini impurity in the fitted random forest (RF) model trained on the dataset shown in A/. G/ Feature importance as measured by endoR and feature influences for each discretized level of numeric variables as computed by endoR. The point color indicates whether features were used to construct the response (‘True’) or not (‘Irrelevant’). H/ Decision network produced by endoR. Edges and nodes correspond to single variables and their interaction effect on the response variable, respectively. Edge widths and node sizes are proportional to the interaction and feature importances calculated by endoR, respectively; their colors are representative of their influence (see Methods and Supplementary Methods, for details on network construction). The edge transparency is inversely proportional to the importance for H only. I/ Same than H but edges with lowest interaction importance were removed to obtain paths between nodes of length *≤* 3. E, G-I/ Levels of discretized variables, i.e., numeric variables transformed into categorical variables based on their quantiles, are shown on the x-axis of the influence plot (G/) and indicated by ‘__High’ or ‘__Low’ in networks (E/ and H/).

**Supplementary Figure 3.**
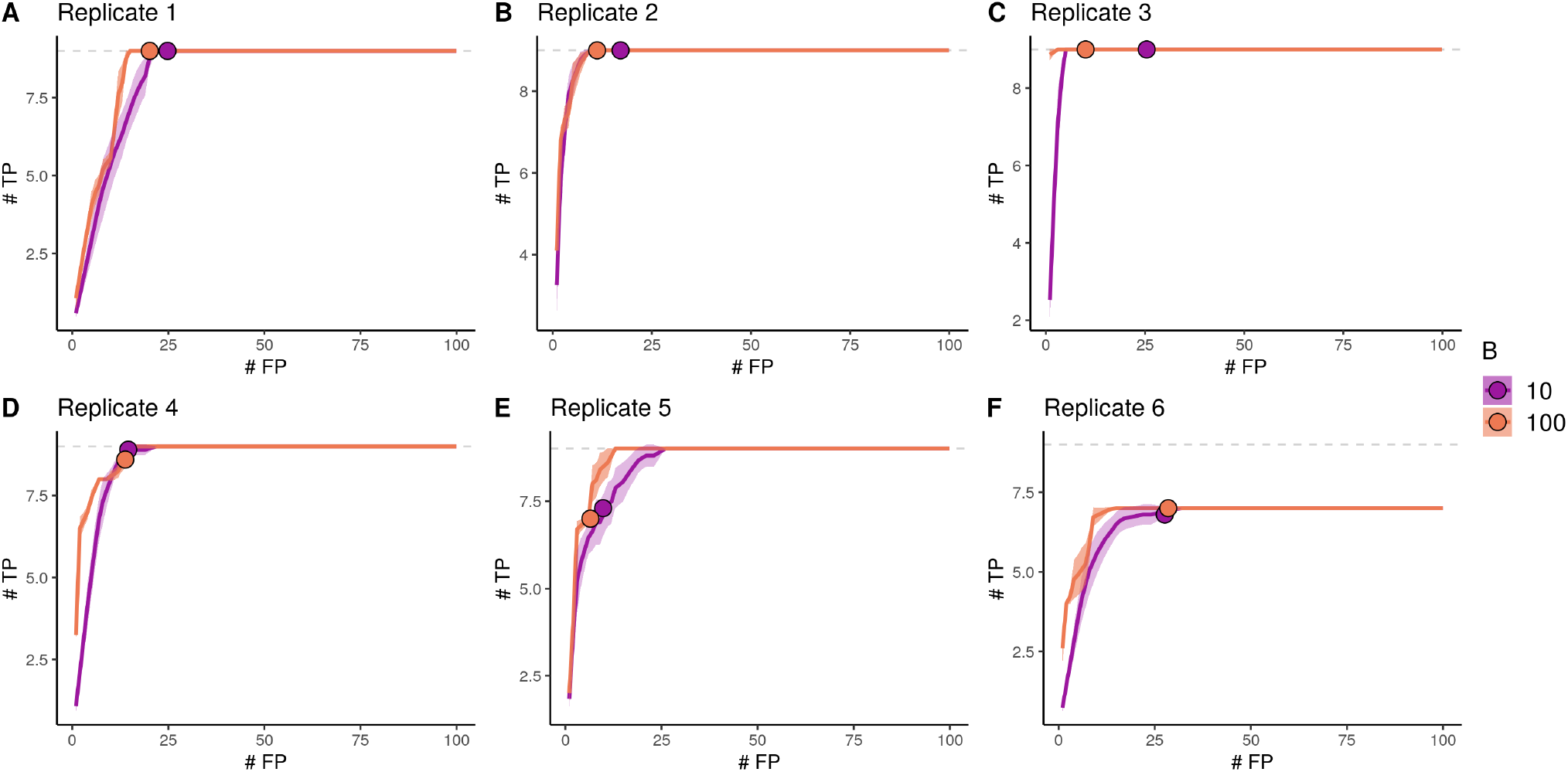
endoR performance stabilizes as the number of bootstraps increases. A total of 6 replicates of artificial phenotypes were each processed 10 times with *B* = 10 or 100 bootstraps resamples (purple and orange, respectively). The curves show the average number (‘#’) of identified true positive (TP) and false positive (FP) edges according to edge probabilities of being selected in the stable decision ensemble. Curves were interpolated for each technical replicate, and the average (line) and standard deviation (shaded area) across number of bootstraps are displayed. The traced points denote average number of TP and FP in the stable ensembles returned by endoR for *π* = 0.7 and *α* = 5.

**Supplementary Figure 4.**
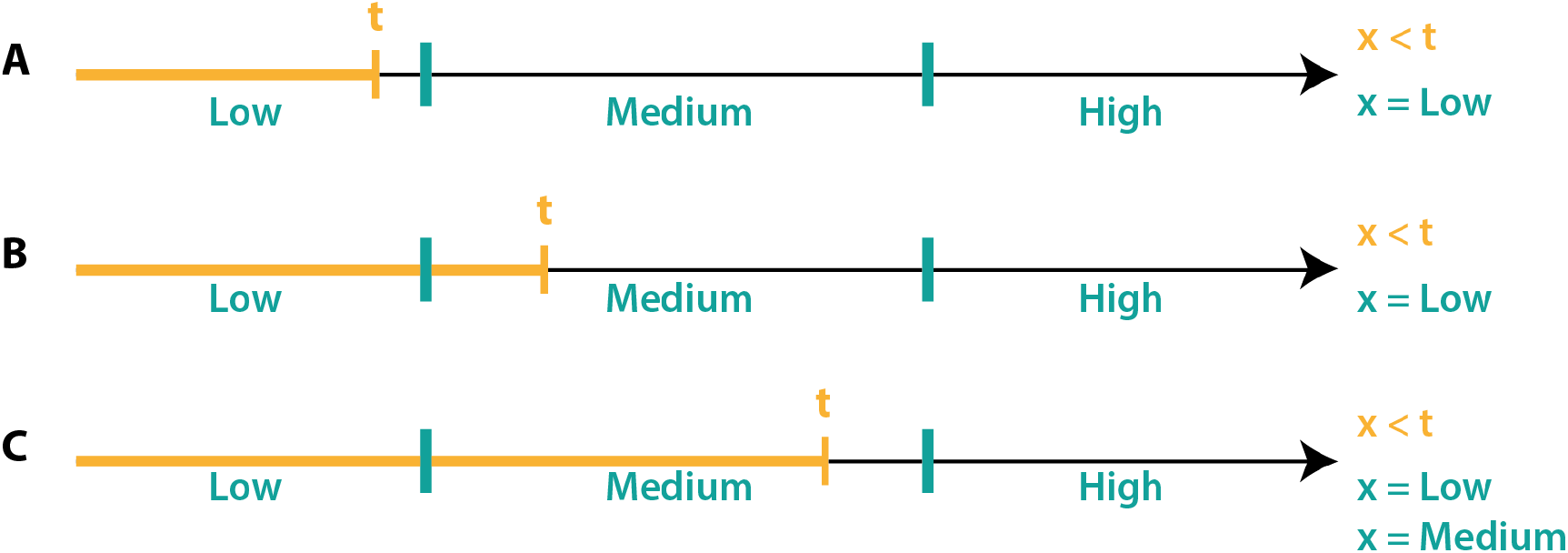
Discretization of variables and modification of rules. Simple example of the discretization of a uniformly distributed variable *x* into three levels. An original rule “*x < t*” (orange) is modified according to the number of observations in each level included in the sample support of the rule (new rule-s in green). B/ A minority of samples in the “Medium” level were included in the original sample support defined by “*x < t*”, therefore the “Medium” level is not selected to make a new rule as in C/.

**Supplementary Figure 5.**
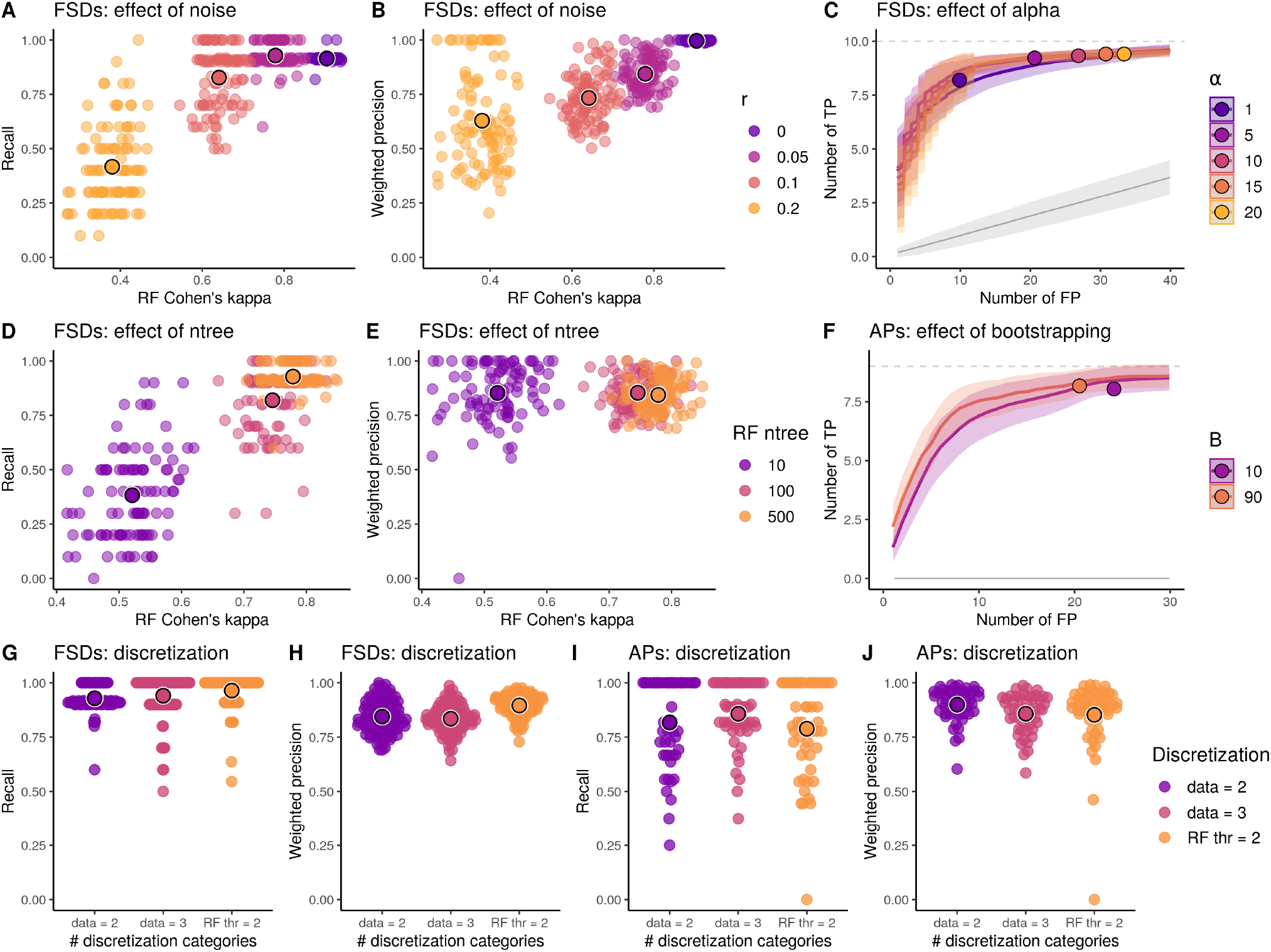
The accuracy of endoR increases with the accuracy of input models. 100 FSDs (B-E and G-H/) and 50 APs (A, F, and I-J/) were generated, RFs were fitted and processed with endoR. If not varied, parameters were as follow: *ntree* = 500, discretization was performed with the method based on the data distribution with *K* = 2 categories, and *α* = 5. We computed the following three metrics: Cohen’s *κ* of the RF, weighted precision and recall values of the selected edges in the stable decision ensemble, and TP/FP-curves based on the probabilities of being selected in the stable decision ensemble (see Methods). A-B, D-E, and G-J/ TP/FP-curves are averaged across all datasets for a fixed parameter setting (line) and standard deviation (shaded area) are displayed. The average number of TPs and FPs expected for a randomization null model and standard deviations, are shown in grey. Large points indicate the average number of TPs and FPs in the stable ensembles generated by endoR. C and F/ Each point corresponds to the precision/recall of endoR applied to a single dataset and parameter setting. The larger traced points are the averages across all datasets for a fixed parameter setting. A-B and D-E/ As expected decreasing the noise or increasing the number of trees in the forest improves the performance of endoR both in terms of precision and recall. Importantly, there is a strong dependence of endoR performance on the performance of the fitted RF and endoR. Moreover, endoR has a good precision even for small RF. C/ Increasing *α* increases both the TPs and FPs. Small values of *α* effectively control the FPs without strongly impacting the recovered TPs. F/ Larger values of *B* are slightly better but endoR performs well even for small values of *B*. G-J/ Discretization was performed by creating K = 2 or 3 categories from numeric variables based on their distribution (‘data’) or on the splits on these variables in the fitted RF (‘RF thr’). Discretization slightly influences endoR performance, without any clear pattern between the FSD and AP simulations.

**Supplementary Figure 6.**
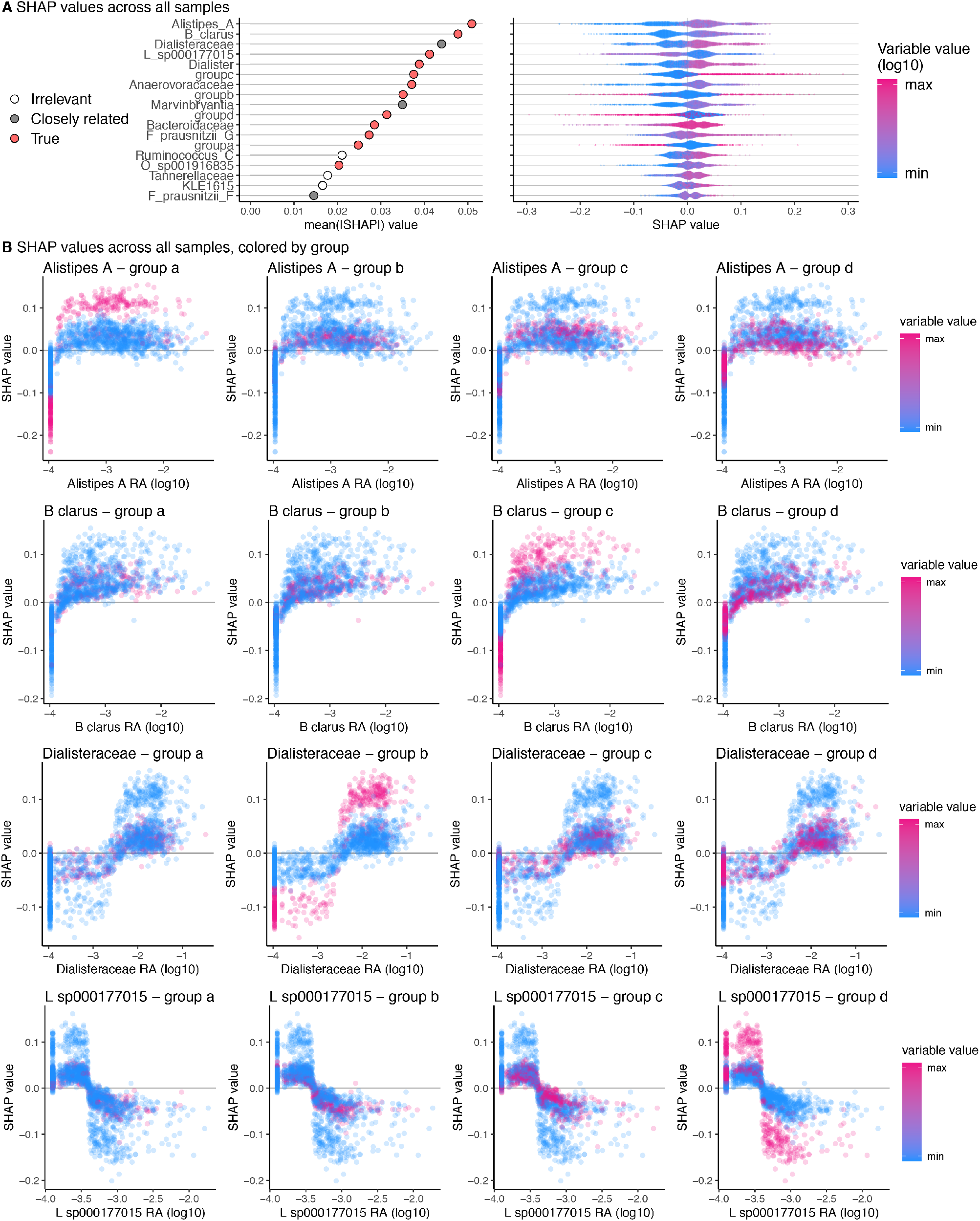
SHAP values from the RF classifier. SHAP values were calculated from the random forest classifier trained to predict an artificial phenotype simulated from real metagenomes (*n* = 2147, *p* = 520 taxa; see Figure 2). A/ The feature importance is given by the average of the absolute SHAP values across samples and is plotted for each sample as well. B/ The SHAP interaction values could not be calculated due to the lack of R-implementation for calculating SHAP interactions from random forests. Consequently, we plotted SHAP values of the four taxa with the highest feature importances (y-axis) according to taxa relative abundances (log10 transformed, x-axis) and colored points by each of the four group category (pink: sample from the group category indicated in the plot title, blue: sample from the other categories). We note that this method of analysis does not scale well as the number of features increases.

**Supplementary Figure 7.**
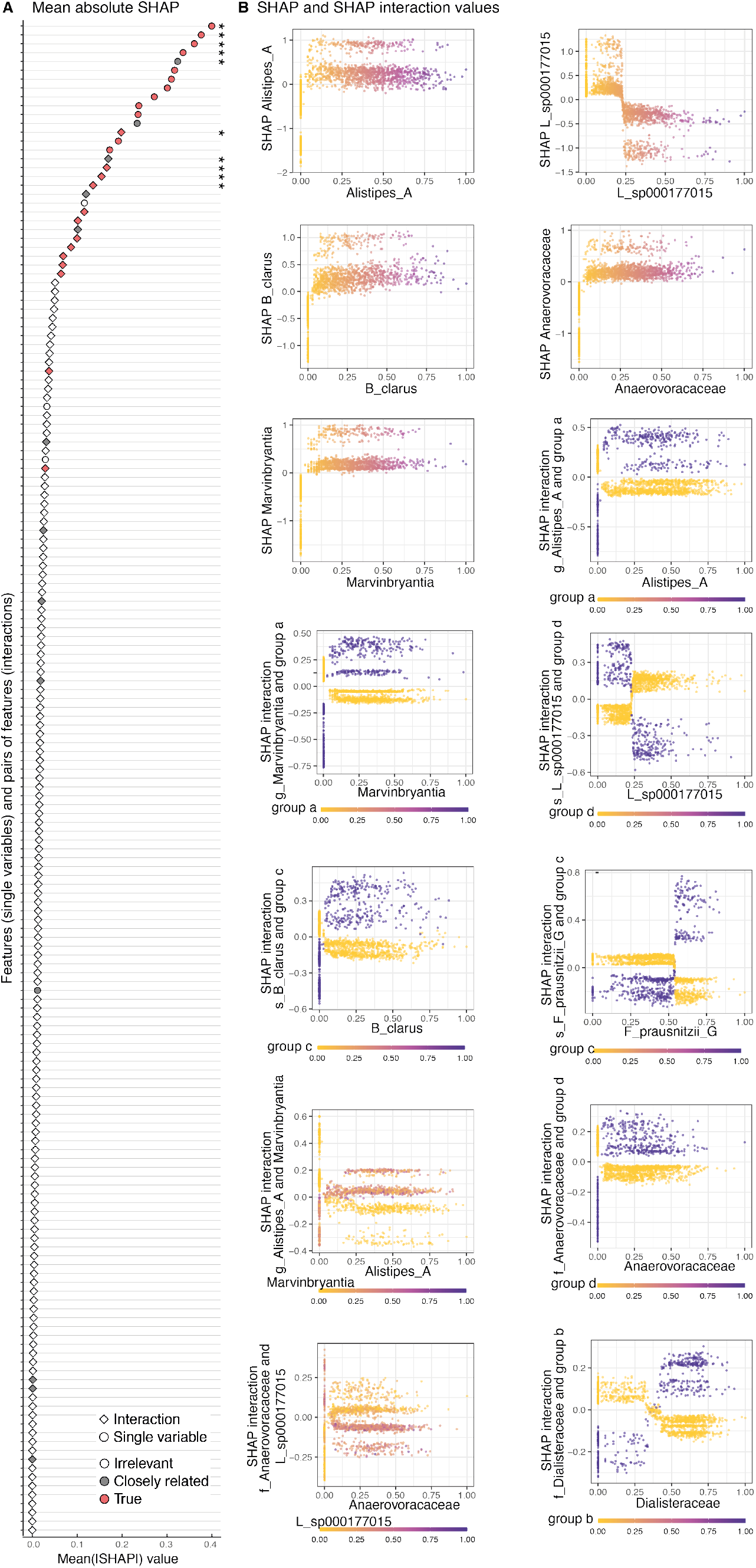
SHAP values from the XGBoost classifier. SHAP values were calculated from the XGBoost classifier trained to predict an artificial phenotype simulated from real metagenomes (*n* = 2147, *p* = 520 taxa; see Figure 2). A/ The feature and interaction importances are given by the average of the absolute SHAP values across samples. B/ Given the high number of features and interactions, we only plotted the top five feature importances of single variables and top nine feature importances for interactions (marked with a start on A/). For single variables, the point color corresponds to the x-axis value.

**Supplementary Figure 8.**
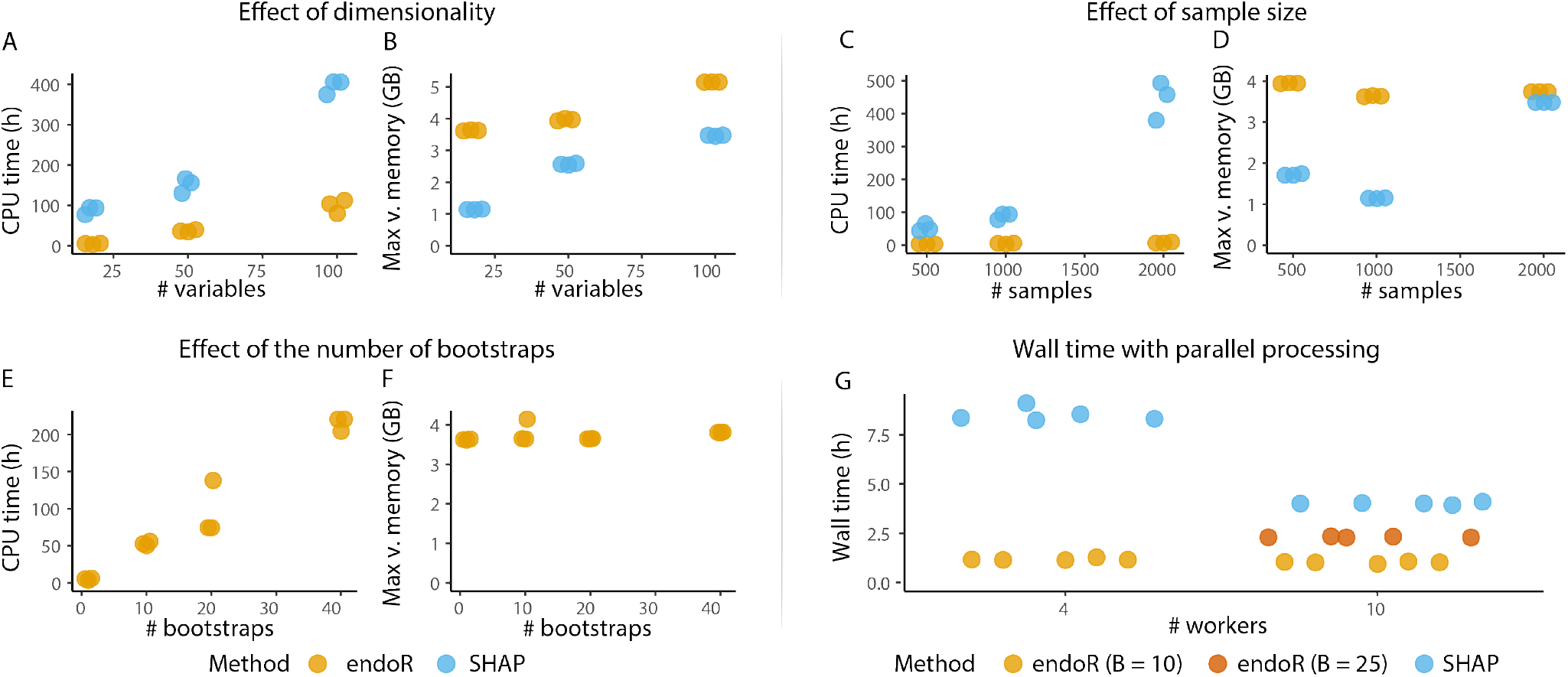
endoR computation time scales substantially better than SHAP when applied to random forest classifiers. A-F/ The total CPU time and maximal virtual memory used for three replicate processing runs of the same RF model. The artificial phenotype presented in Figure 2 was used with 18 variables and 1000 samples (see Methods), and endoR was run on *B* = 1 bootstrap of size *n/*2. G/ Five technical replicates of endoR and `shap` runs on the RF trained to predict the artificial phenotype presented in Figure 2 (18 variables and 2147 observations). Calculations were ran in parallel across 4 or 10 workers; for endoR, bootstraps were further ran individually in parallel (see Supplementary Methods).

**Supplementary Figure 9.**
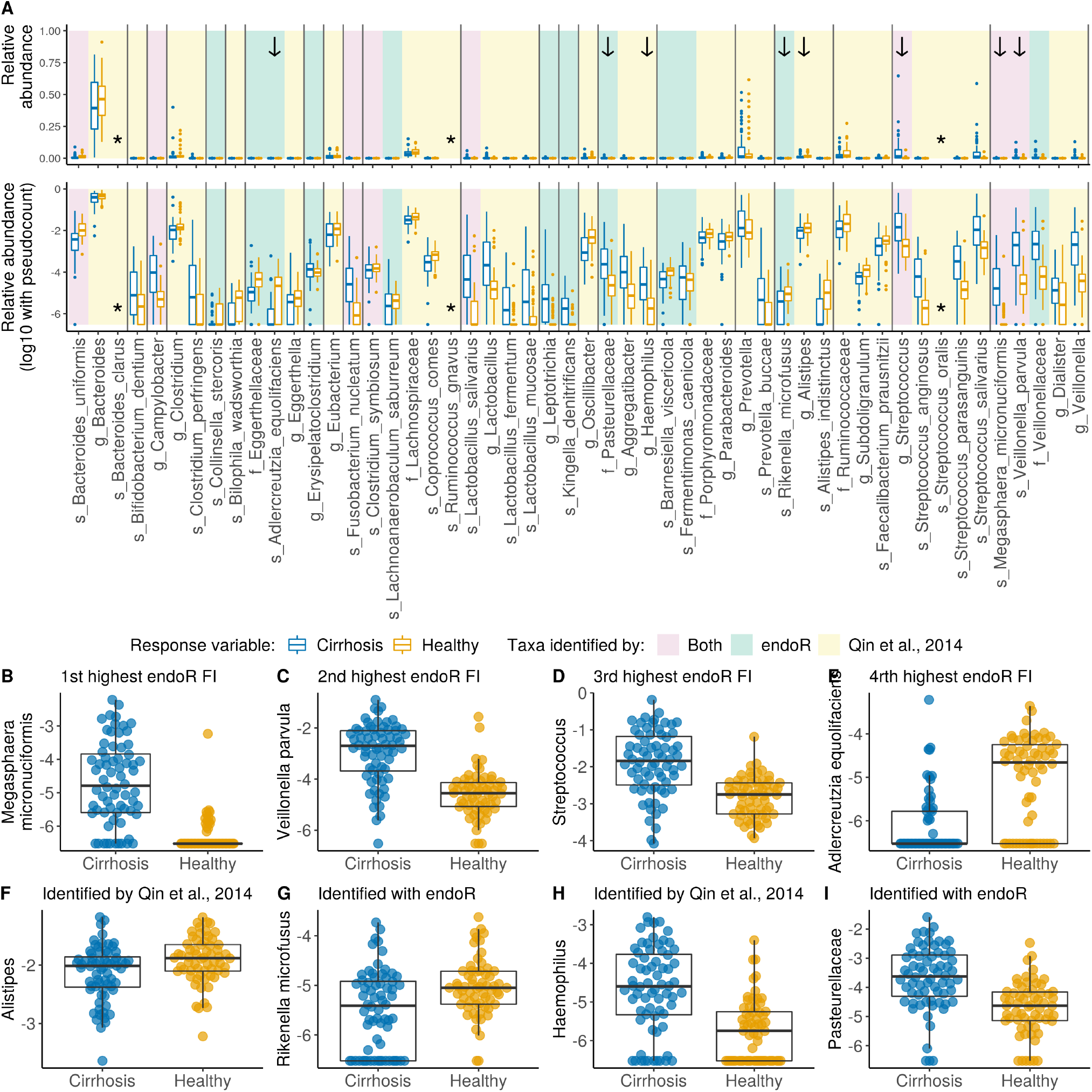
Relative abundances of taxa identified using a RF and endoR versus statistical tests in the original study (Qin et al., 2014). When the log10 of relative abundances is displayed, a pseudo-count equal to the minimal relative abundance detected in the dataset (3 *·* 10^−7^) was used to show samples for which taxa were not detected (relative abundance = 0). Boxplots and points are colored by healthy status, with healthy individuals in orange and cirrhotic ones in blue. A/ Taxonomic levels are indicated with the prefixes: ‘f_’ = family, ‘g_’ = genus, ‘s_’ = species. Taxa are organized by family taxonomic level (separated by grey lines). The background indicates whether taxa were identified in this article and the original study (red), only in this article via a RF model and endoR (green), or only in the original study (yellow). Species for which the relative abundances were not available in the published dataset (downloaded from the ML task repository) are indicated with a star. Taxa in B-I/ are indicated by an arrow. B-E/ The four taxa taxa with highest feature importance (FI) identified by endoR to classify healthy versus cirrhotic microbiomes (see Figure 5 A). F-I/ Taxa exclusively identified in the original study (Qin et al., 2014) or with a random forest and endoR.

**Supplementary Figure 10.**
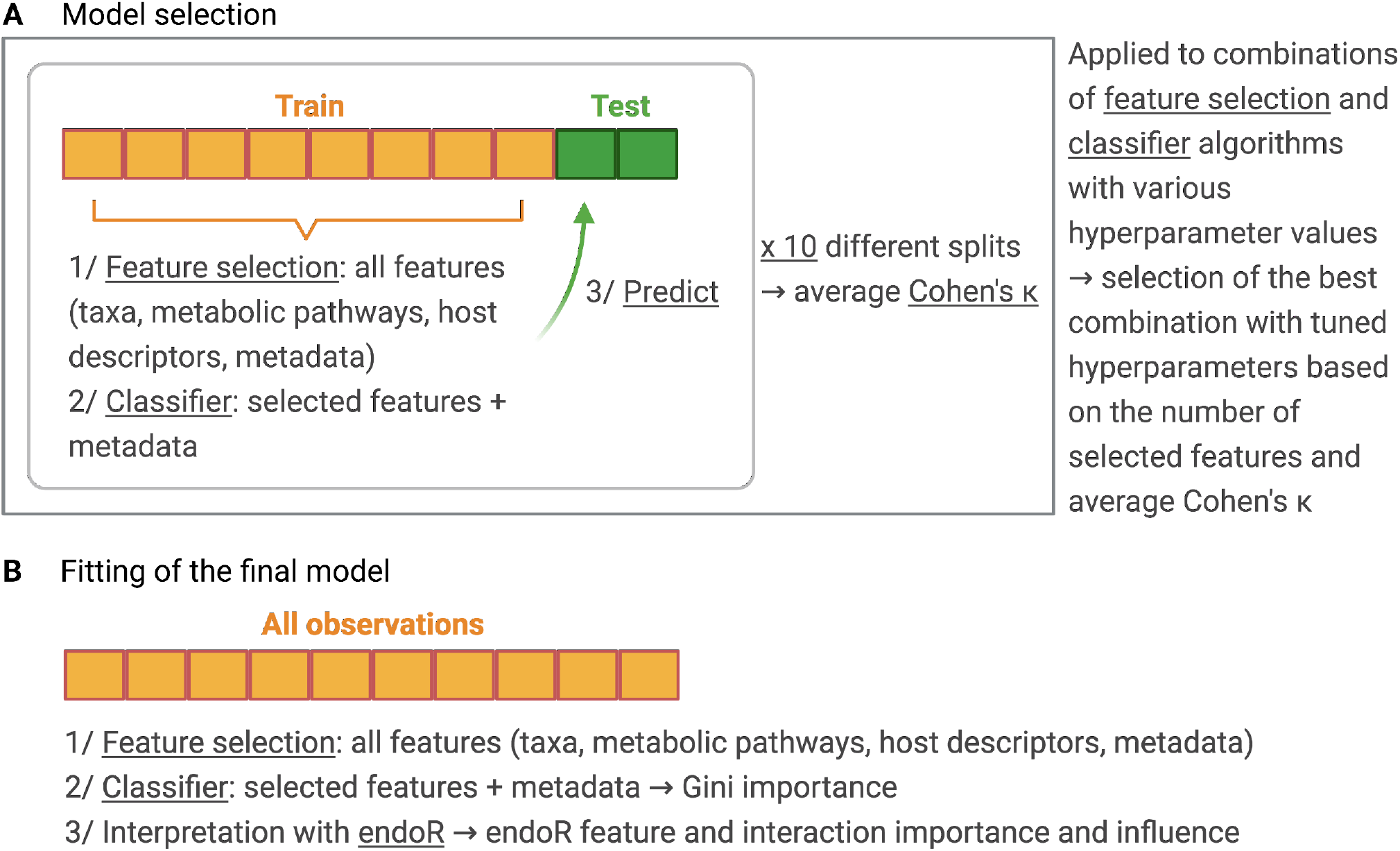
Model selection and fitting for predicting the presence/absence of *Methanobacteriaceae*. A/ Ten sets of observations, each containing a subset for training and one for testing, were created. Training observations were used to fit models, i.e., the combination of a feature selection and classifier algorithms for given hyperparameter values, and testing ones were predicted with the fitted models. Model’s performances were averaged across testing sets. Feature selection algorithms consisted of (i) no feature selection, (ii) a taxa-aware version of the gRRF algorithm from Deng (2013) (Supplementary Methods), (iii) the Boruta algorithm (Kursa et al., 2010), (iv) no feature selection. Classifiers were fitted with random forests or gradient boosted model algorithms. Metadata correspond to the number of reads and original dataset names. B/ The model (feature selection algorithm and classifier) that resulted in A/ in the highest average Cohen’s *κ* using the fewest features was used to fit the final classifier on all data.

**Supplementary Figure 11.**
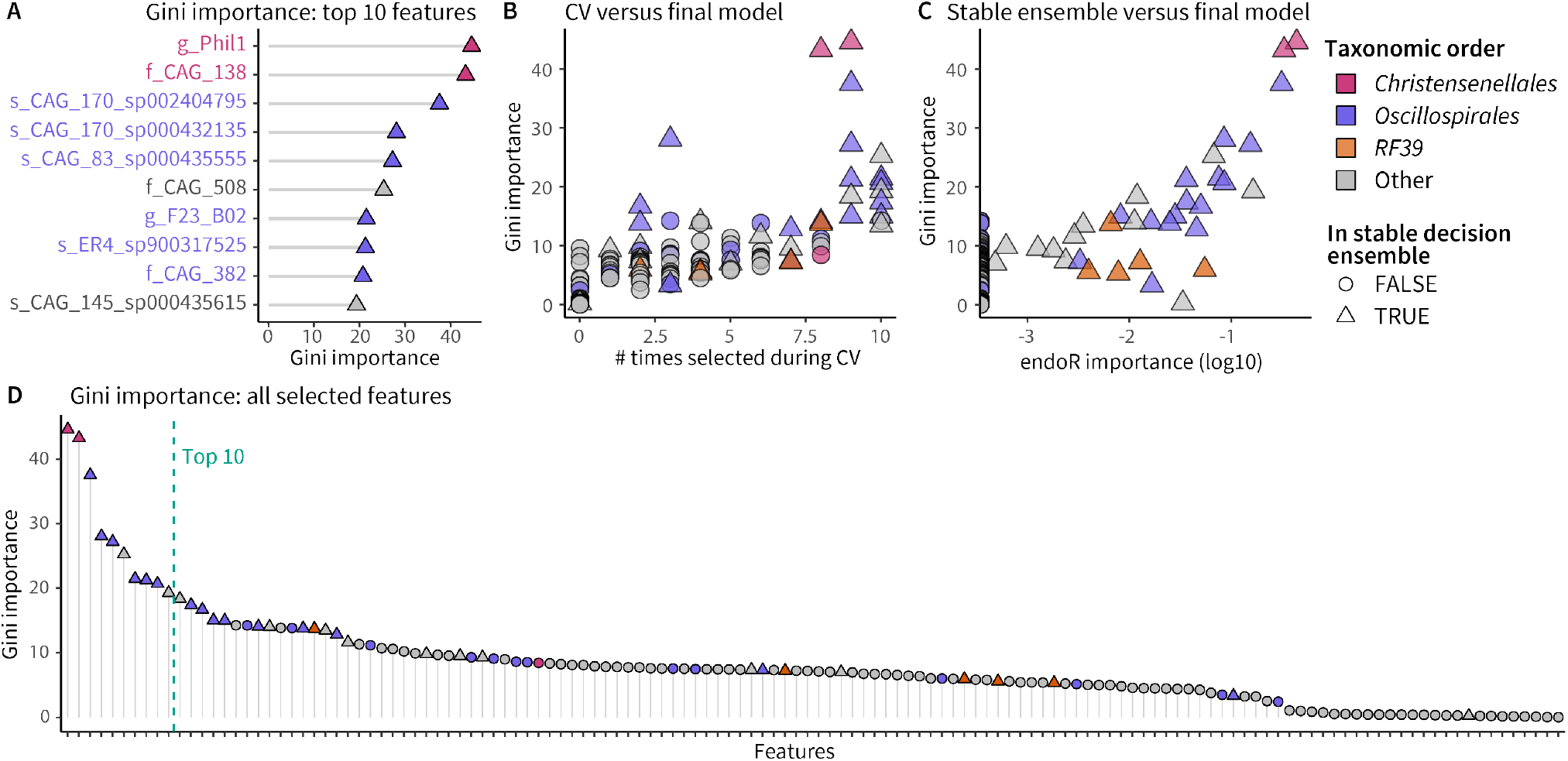
The Gini and endoR importances are consistent between the RF model and stable decision ensemble. A/ Features with the best Gini importance. B/ Comparison of the Gini importance and the number of times features were selected across the 10 cross-validation (CV) sets. C/ Comparison of the Gini and endoR importance. D/ Gini importance of all features selected by the taxa-aware gRRF algorithm. In all plots taxonomic levels are indicated in the labels with ‘f_’: family, ‘g_’: genus, and ‘s_’: species, and orders are indicated via point and label colors.

**Supplementary Figure 12.**
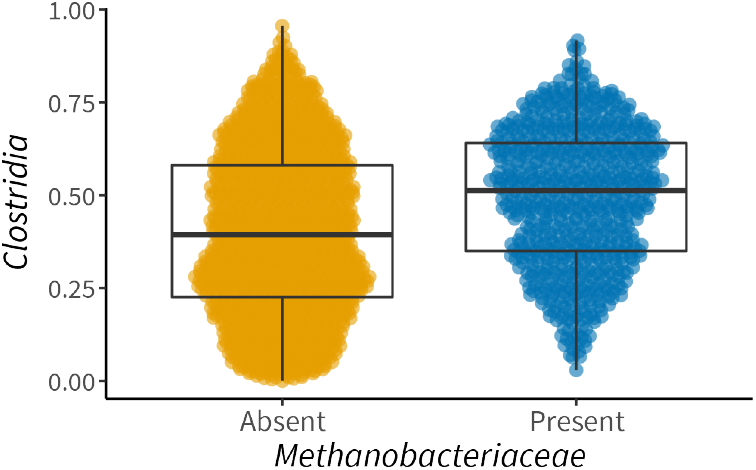
The relative abundance of *Clostridia* is higher in samples where *Methanobacteriaceae* are detected.

**Supplementary Figure 13.**
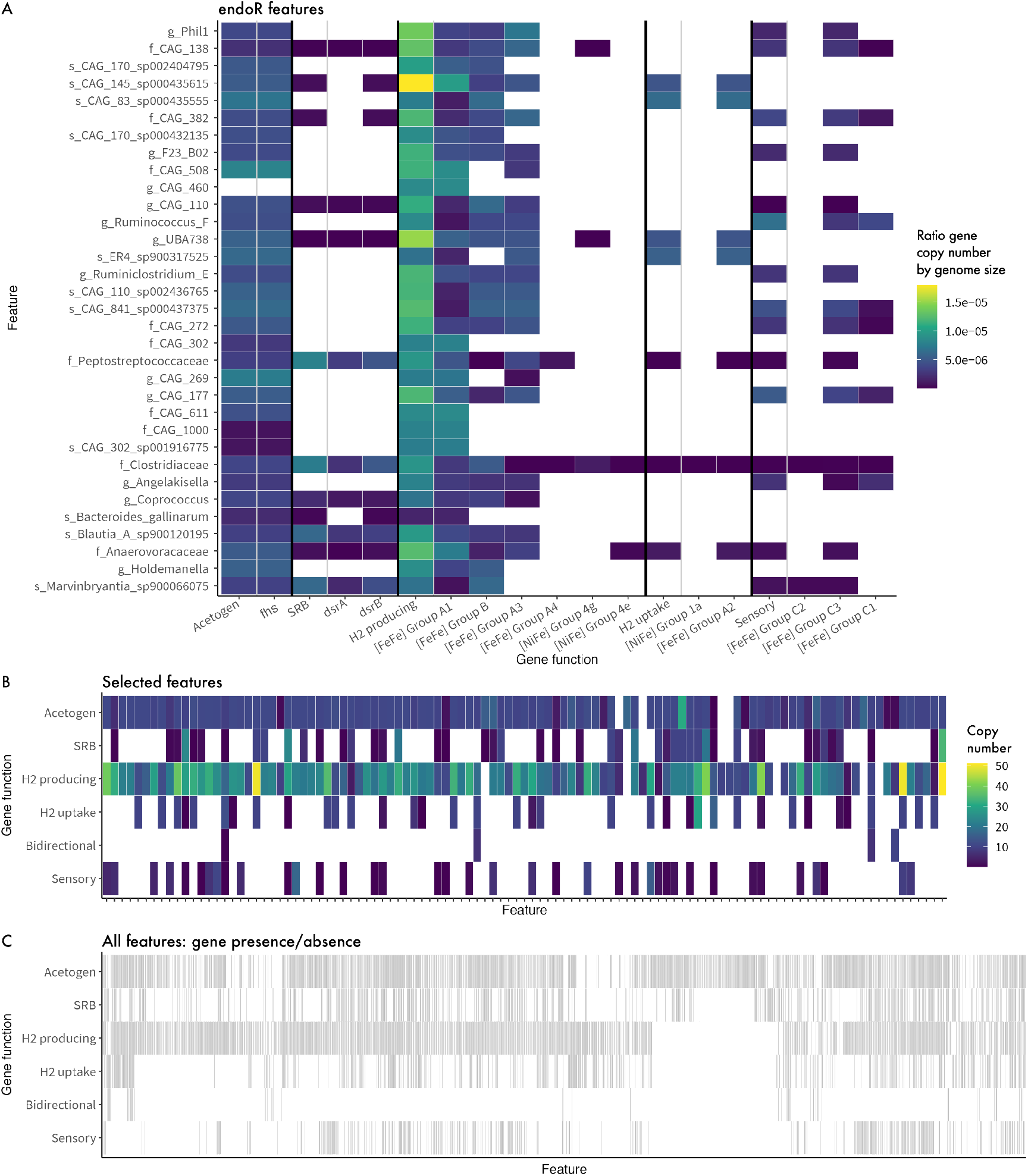
Copy number of genes involved in H_2_ consumption and production across taxa used to predict the presence/absence of *Methanobacteriaceae*. We looked into the genomes of representative species of taxa used to predict the presence/absence of *Methanobacteriaceae* in human guts microbiome from 2203 individuals for involved in H_2_ metabolism. The number of copies of genes involved in the following pathways or function were counted: sulfate reduction (SRB): *dsrA* and *dsrB* genes (Fish et al., 2013); acetogenesis (Acetogen): *fhs* gene (Singh et al., 2019); H_2_ production, uptake and sensing as determined by the HydDB database (Søndergaard et al., 2016). At the genus and family taxonomic levels, we used the average number of copies across species from the given level and weighted the number of copies of each species by the average relative abundance of species in the dataset. Accordingly, if the most abundant species of a specific genus had high number of gene copies, the number of copies for that genus would also be high. When genes were grouped by general function, we summed the number of copies (e.g., the SRB gene copy number corresponds to the sum of gene copies of *dsrA* and *dsrB*). A/ The ratios of gene copy number by genome size for each of the endoR selected features are consistent with the absolute number of copies displayed in Figure 6 B. For each representative species, the number of gene copies was divided by the genome size. General functions and genes are displayed and separated by black lines (blocks of genes with the same general function), general functions are separated from specific genes by a grey line. B/ Number of genes copies from each group for taxa selected by feature selection. C/ Occurrence of genes from each group across all taxonomic features used to train models to predict the occurrence of *Methanobacteriaceae* in human guts.

**Supplementary Figure 14.**
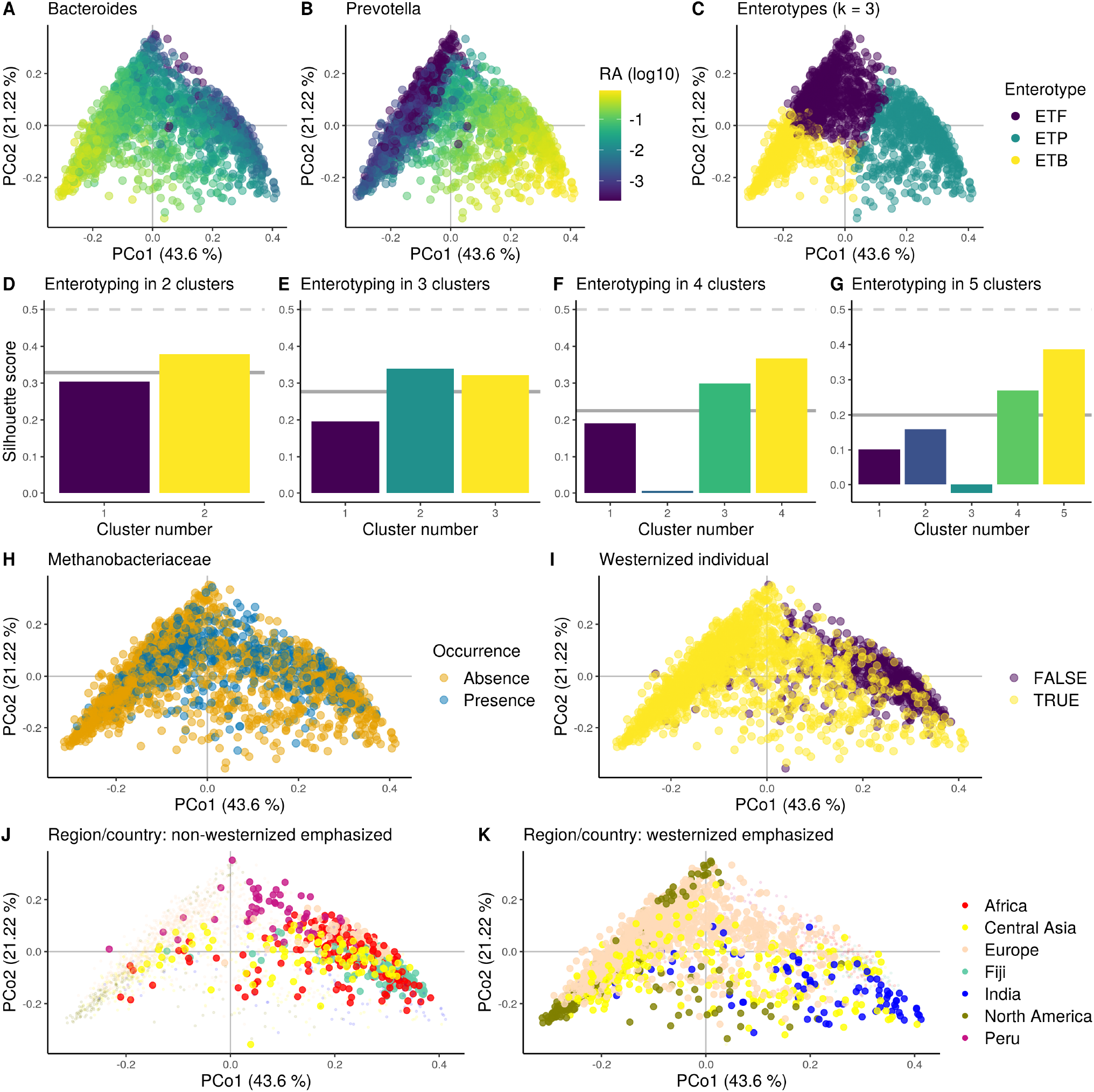
Gut microbiota of a large cohort of healthy individuals (n = 2203 individuals) approximately segregate along the enterotype landscape. A-C, H-K/ Principal coordinate analysis ordination of the Jensen-Shannon distance matrix computed from genera relative abundances. A-B/ Samples colored by relative abundance (RA) of *Bacteroides* and *Prevotella*, respectively. To calculate the log, a pseudocount equal to the minimal non-null RA was given to samples for which the genus was not detected, i.e., RA = 0. C/ Samples colored by enterotype cluster (Arumugam et al., 2011; Costea et al., 2018); ETF: *Firmicutes*, ETB: *Bacteroides*, ETP: *Prevotella*; colors correspond to those on E. J-K/ Samples are colored by their country of origin grouped by region when possible (e.g., Canada and USA grouped into North America; Supplementary Table 7), and points are emphasized (larger and less transparent) if they were sampled from non-westernized (J) or westernized (K) populations. D-G/ Average silhouette score (bar) within each k-mean cluster computed from the Jensen-Shannon distance matrix and across clusters (thick line). Dashed line: threshold above which clustering strength is moderate (Koren et al., 2013).

**Supplementary Figure 15.**
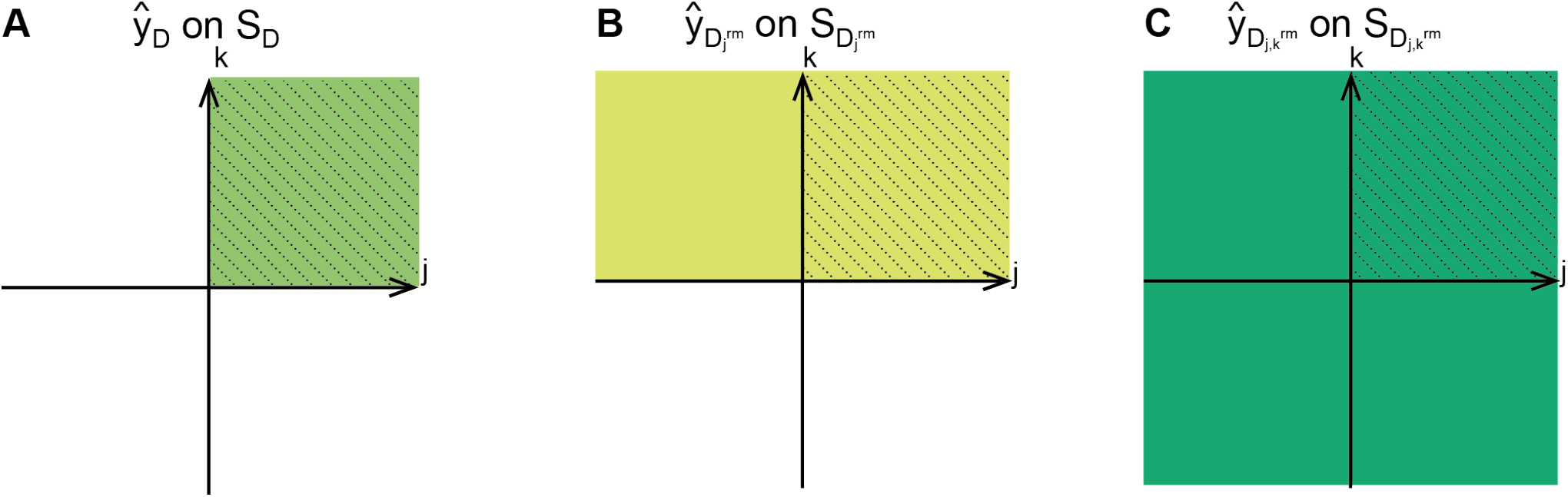
Visualization of how decisions are modified to calculate the importance of variables. Each plot illustrates the support of a decision *D* in the feature space spanned by variables {*x*^*j*^, *x*^*k*^}, i.e., the values that the decision can take on variables *x*^*j*^ and *x*^*k*^. A/ Original decision *D*. B/ Modified decision 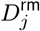 resulting from removing variable *x*^*j*^ from decision *D*. C/ Modified decision 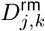 resulting from removing variables *x*^*j*^ and *x*^*k*^ from decision *D*. A-C/ The support *S*_*D*_ of the originl decision is indicated by the stripped areas, such as samples in the support of *D* all take positive values on *x*^*j*^ and *x*^*k*^. The support of each decision, i.e., *S*_*D*_, 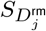 and 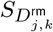 for A, B, and C, respectively, is visualized by the colored region. B/ When we remove variable *x*^*j*^ from the rule *r*_*D*_ of *D*, the support 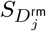 is extended to samples taking negative values on *x*^*j*^ (colored area). C/ Similarly, when we remove a pair of variables *{x*^*j*^, *x*^*k*^ *}* from *r*_*D*_, samples in 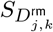 can take positive and negative values on *j* and *k*. For 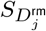 and 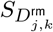, we calculate 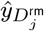 and 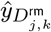, respectively, using all samples in 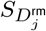 and 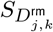. The decision-wise importance 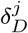 of *j* in *D* is calculated by comparing the error of 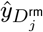 on *S*_*D*_ (B/) versus the error of *ŷ*_*D*_ on *S*_*D*_ (A/). Similarly, to calculate the decision-wise importance of a pair of variables {*j, k*} in a decision *D*, we compare the error from the decision not constraining values on *j* or *k*, with 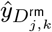 on *S*_*D*_ (C/) to the error of the decision with *ŷ*_*D*_ on *S*_*D*_ (A/).

**Supplementary Figure 16.**
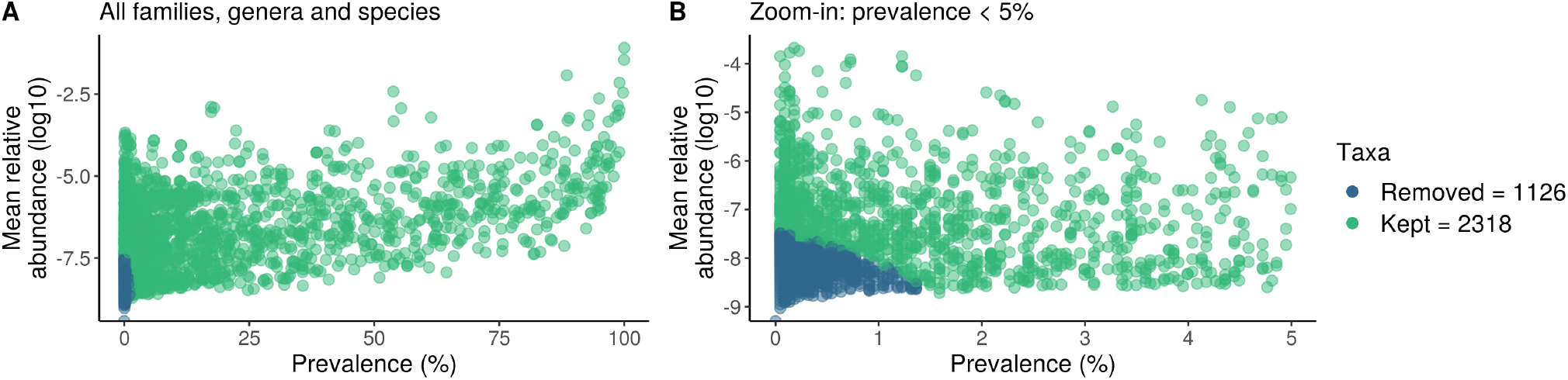
Mean relative abundances and prevalence of family, genus and species taxonomic levels in the metagenomic data.

## Notes

https://github.com/leylabmpi/endoR

https://github.com/aruaud/endoR_data_analysis

### Competing Interest Statement

The authors have declared no competing interest.

